# The reliability of environmental cues shape learning and selection against deleterious alleles in seed beetles

**DOI:** 10.1101/2025.03.23.644788

**Authors:** Lynne Caddy, Tessy Munoz, Julian Baur, Loke von Schmalensee, David Berger

## Abstract

Behavioural plasticity can play a key role in evolution by either facilitating or impeding genetic adaptation. The latter occurs when behaviours mitigate selection pressures that otherwise would target associated traits. Therefore, environments that facilitate adaptive behavioural plasticity could relax the strength of natural selection, but experimental evidence for this prediction remains scarce. Here, we first demonstrate that maternal care in the beetle *Callosobruchus maculatus* is dependent on environmental cues that allow females to reduce larval competition via learning and informed oviposition choices. We show that this facilitation of maternal care relaxes selection against deleterious alleles in offspring. We further find that mothers of low genetic quality generally provide poorer care. However, when receiving environmental cues providing accurate information about future host-quality, the increased opportunity for adaptive behavioural plasticity reduced genetic differences in maternal care, further relaxing selection against deleterious alleles. We use our data to illustrate how the identified link between adaptive behavioural plasticity in maternal care and the strength of natural selection can impact indirect genetic effects between mothers and offspring and the accumulation of cryptic genetic loads in populations inhabiting environments that differ in their predictability.

## Introduction

Phenotypic plasticity plays a fundamental role in shaping organismal responses to changing environments (1–4). Plasticity can help organism to survive and reproduce despite initial adversities. This can allow populations to persist in novel environments and respond to selection, leading to increased genetic change over time (1,2,5,6). However, by mitigating selection pressures, phenotypic plasticity can also reduce genetic adaptation (2,7,8). This forgiving nature of plasticity should relax purifying selection and lead to the accumulation of genetic variation that would be deleterious in the absence of plasticity (9–11). If this process continues over time, a population could become dependent on plasticity to survive, as without it, the built up load of conditionally deleterious alleles would become exposed to selection (10,12–14).

The evolution of adaptive plasticity is contingent on the organism’s ability to change phenotype in accord with changes in the environment (9,15,16). This leads to the expectation that environments that provide reliable information of future change should favour the evolution of adaptive plasticity, whereas unpredictable environments may hinder it when environmental fluctuations are too fast, and/or costs of plasticity are too high, to allow accurate phenotype-environment matching (9,15,17,18). Many behavioural phenotypes represent striking forms of plasticity, reflecting an organisms’ use and processing of information through experience and learning to fine-tune its strategies to fit the perceived environment (3,19,20). Indeed, adaptive behavioural plasticity in the form of learning has been argued to be a key determinant of evolution in novel and fluctuating environments (2,3,9,21,22). Like for other forms of plasticity, however, learning will only be selected for if reliable environmental cues are available, allowing for behavioural adjustments that accurately match future conditions (9,15,23). This suggests that populations inhabiting environments that differ in their predictability should exhibit systematic differences in learning and genetic load in associated traits. These dynamics are expected to have important consequences for both the levels of segregating genetic variation in natural populations and for predicting responses to future environmental change (1–3,6,9,16,24,25).

Here, we investigated how the reliability of environmental information impacts the opportunity for adaptive behavioural plasticity and the resulting strength of selection on deleterious genetic variation in the seed beetle *Callosobruchus maculatus* (**Fig. 1**). We studied behavioural plasticity in maternal care and females’ ability to use information to assess future egg-laying opportunities and learn the spatial location of high-quality hosts. The immediate survival benefits of maternal care to offspring are obvious (26,27), but the flip-side of the coin is that by relaxing selection in offspring, maternal care should, at least theoretically, allow deleterious genetic variation to accumulate. Indeed, in burying beetles, populations that are deprived of parental care suffer immediate increases in juvenile mortality and inbreeding depression (28,29), but at the same time seem to more efficiently purge genetic variation in the long run (11,12,30). Moreover, given that cognitive processes are costly and governed by many genes with pleiotropic effects (9,31–33), the build-up of genetic load via relaxed selection in offspring could feed back on parents’ ability to learn and provide care. Genetic variation in parental care is well-known to impact the expression of genetic variation and selection on traits in offspring via indirect genetic effects (*IGEs*: the effect of genetic variation expressed in one individual on trait expression in interacting individuals (34–36)). Imagine that parents of high genetic quality provide better care for their young, who also have intrinsically higher survival due to their superior genetic quality. This positive covariance between parent and offspring breeding values increases genetic variance in offspring survival and is predicted to speed up responses to selection compared to a situation where IGEs are absent (34,36,37). Thus, if the reliability of environmental information affects the amount of expressed covariance between adult and juvenile traits, it could have strong effects on evolutionary dynamics (14,24,38).

**Figure 1:**
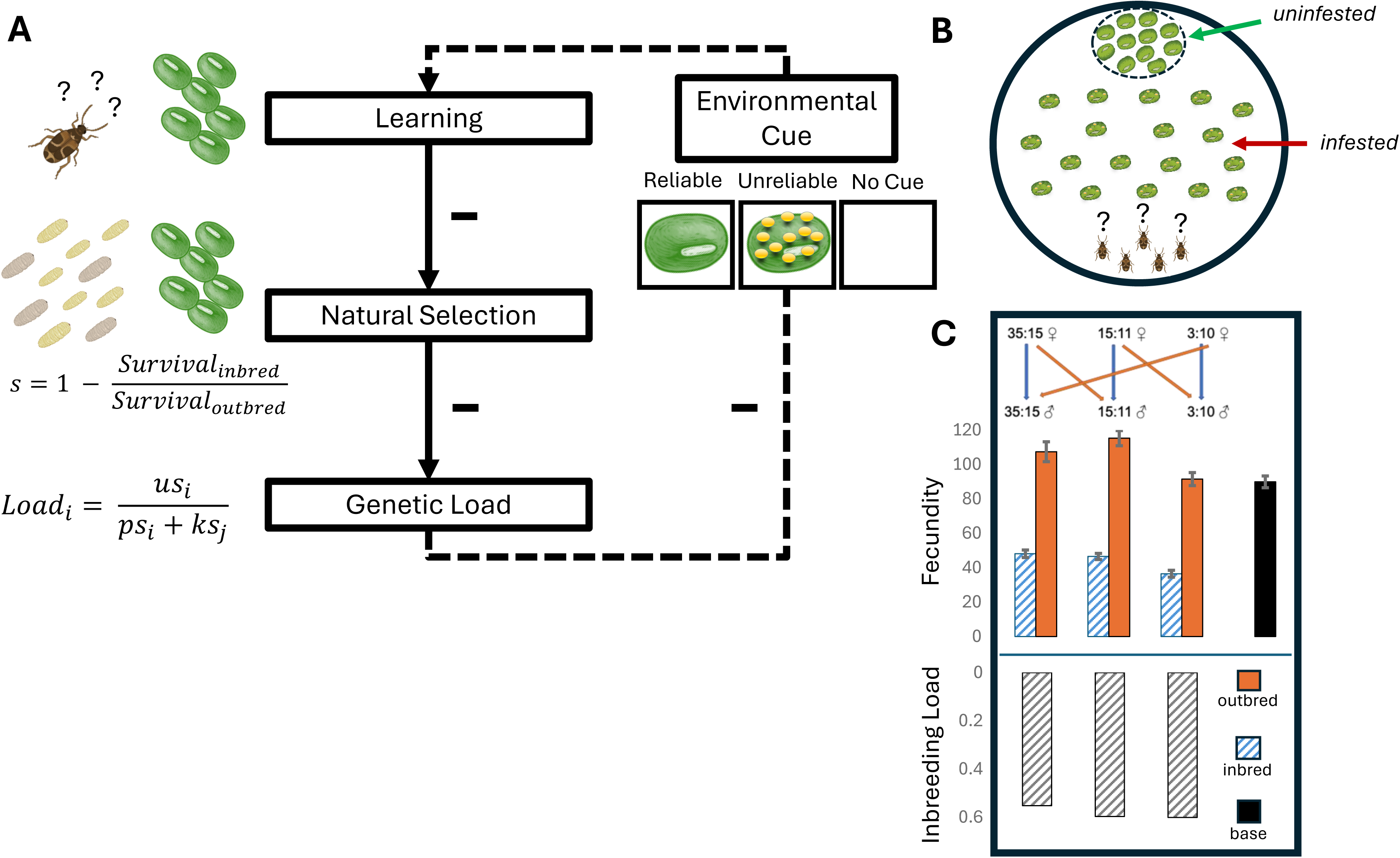
Study design. **A)** Hypothesized relationships between the reliability of environmental cues, learning, natural selection, and genetic load. Reliable environmental information is expected to favour adaptive behavioural plasticity in form of learning. Behavioural plasticity can mitigate genetic differences in trait expression and relax the strength of natural selection, thereby increasing the build-up of deleterious genetic variation over time (i.e. genetic load). The accumulation of genetic load could in turn impair learning (broken arrow). However, the degree to which could depend on the available environmental cues used in the learning response. We assessed how environmental cues affected adaptive behavioural plasticity in maternal care, and how this in turn affected selection in larval offspring. **B)** Female *C. maculatus* were challenged to find and discriminate among hosts seeds in a spatial learning task. The seeds were either uninfested or infested with 2-4 eggs of the competitor *C. phaseoli.* We gave females three different treatments before they performed the learning task; a reliable cue (uninfested seeds) giving accurate information about the presence of future high-quality hosts, an unreliable cue giving false information (seeds infested with >10 eggs), or no cue at all. We measured the fitness consequence of maternal care behaviour and learning in each treatment by combining data on female oviposition choice with estimates of larval survival. **C)** Three highly inbred strains (blue stripes) were used and compared to themselves when outcrossed (orange) (base stock population; black). Fecundity estimates per line are given as mean ± 2 se. The inbred strains show inbreeding depression amounting to a reduction in fecundity of 55-60% relative to when outcrossed (grey striped bars). The comparison between the three strains in inbred and outbred state was used to assess the relative strength of viability selection in larvae (*s*) and the effect of recessive genetic load on female learning and maternal care, across the three treatments. Based on these estimates, we predicted the genetic load at mutation-selection balance in a meta-population where individuals received reliable information, *i*, or no information, *j*, at fraction *p* and *k* as a function of the deleterious mutation rate, *u*, and the strength of selection, *s*. See main text for further details.

Female seed beetles provide care for their offspring by preferentially placing eggs on uninfested host seeds, a behaviour that can be fine-tuned by experience (39–41). First, we gave females the opportunity to choose between uninfested and infested hosts in a spatially complex environment that they were exposed to multiple times. Before and in-between exposures, they received either reliable or unreliable information about the environment’s host infestation rate. We show that learning of the spatial location on high-quality hosts is faster, and care behaviour more accurate, if females receive accurate information about potential future hosts. Second, we find that viability selection in offspring is significantly reduced, and juvenile survival increased, by improved care provided by mothers receiving accurate information. Third, by comparing care behaviour in mothers of different genetic quality, we find that the improved learning facilitated by accurate environmental information also reduces selection on deleterious alleles in adults. Together, this rendered a 30% reduction in the strength of natural selection and substantially weakened IGEs between maternal care and offspring survival in the treatment providing accurate environmental information. Finally, we adopt an existing theoretical model and parametrize it with our experimental data to illustrate potential population-level consequences of the evolutionary feedback between adaptive behavioural plasticity, the strength of natural selection, and the accumulation of conditionally deleterious genetic variation. The results imply that populations evolving exclusively in predictable environments can build up substantial cryptic genetic loads that would make them inferior competitors in unpredictable environments where adaptive behavioural plasticity is restricted.

## Methods

### Study species

*Callosobruchus maculatus* is a capital breeding beetle pest of leguminous crops in tropical and subtropical regions. Adults do not require food or water to reproduce (42). Adults typically die 7-14 days after emergence in the absence of food and water in laboratory conditions of 29°C and 50%RH (43). Both sexes can start reproducing on the day of adult eclosion and females lay 70-90% of their eggs during the first few days of reproduction (42). The juvenile phase is completed in 3-4 weeks, and egg-to adult-survival rate on *Vigna unguiculata* (black eyed bean) and *V. radiata* (mung bean) is around 90% in laboratory conditions (44–46). Female host search and discrimination is complex. *C. maculatus* uses a wide repertoire of fabaceus host plants (47) but discriminate between high and low quality species (48,49). Females also heavily discriminate against egg-laden seeds and actively avoid competition with other females (39), as high larval density reduces survival and size at maturity (49–51). *C. maculatus* utilizes both natural habitats where host plant patches are widely distributed, as well as grain storages where adult food is absent but egg laying substrate is abundant and population density is higher (47,49,50).

#### Isogenic lines

The three genetic lines used in the experiments were derived from *Vigna unguiculata* (black-eyed bean) seed pods collected at a small-scale agricultural field close to Lomé, Togo (06°10’N 01°13’E) in October and November, 2010 (52). Several lineages were exposed to successive full-sib matings (53), and the three most productive lineages used in this study were continued for 30 generations, resulting in near isogenic strains. Their near complete homozygosity should have prevented further evolution of the strains during the experiments and limits within-strain variation due to segregation. We reasoned that the comparisons of these strains in inbred and outcrossed state would serve as a powerful tool to inspect the fitness consequences of genome-wide homozygosity and estimate the cumulative strength of selection across genomic sites harboring deleterious alleles. Although inbreeding depression can result from increased homozygosity at sites harboring rare weakly deleterious (partially) recessive alleles, as well as at sites harboring intermediate frequency polymorphisms maintain under balancing selection, the former mechanism seems to contribute most to inbreeding depression in natural populations (54), which is also consistent with data from this panel of inbred lines (55). Hence, differences between inbred and outcrossed strains in our study should mainly reflect deleterious recessives, while effects at sites under balancing selection are expected to have a more minor contribution but should not be ruled out completely.

To provide an estimate of the inbreeding load in the three lines, we measured fecundity (number of hatched eggs produced over the first week of reproduction) in females that were either produced by within-(inbred) or between-(outbred) line-crosses. Outcrossing between the three lines was performed in a round-robin fashion following a particular schedule (**Fig. 1C**) and should hide the fitness effects of segregating recessive deleterious alleles, given that these are relatively rare and unlikely to be shared among the three lines. To ensure that bad sperm from inbred males played no part in the fecundity estimates, all females were mated with males from the outbred Lome stock population. We assayed 15 females from each of the six crosses (three inbred and three outcrossed).

For comparison we also scored the fecundity of 15 females from the Lome stock population. Each female was placed into a 35mm petri dish filled with mung beans. These petri dishes were placed in a climate cabinet set at standard conditions and the females were left to lay for the next 7 days after which females were removed. After another 7 days, beans were removed and the number of hatched eggs were counted. This showed that inbreeding has led to a decrease in female fecundity of 55-60% relative to the outbred state (**Fig. 1C**, Supplementary 1).

#### Larval competition

To estimate the fitness effects of variation in maternal care of *C. maculatus*, we let females choose between uninfested mung beans and mung beans infested with larvae of the closely related competitor, *C. phaseoli*, which has strongly overlapping geographical distribution and host plant preference with *C. maculatus* (56,57). The two species also have similar life-histories (58), although the lab stock of *C. phaseoli* has a 20-30% smaller body size than *C. maculatus* (59). The relative success during interspecific competition inside host seeds could be quantified by counts of offspring emerging from beans as the species differ in colour patterning. All experiments were performed using mung beans, which is a preferred host of both species but small enough to typically allow for only one or two adults to emerge (**Fig. 1A**).

### Learning and maternal care with and without reliable environmental cues

Based on the previous findings (41), we constructed arenas from 150 mm diameter petri dishes. Each arena consisted of a ‘starting zone’ at one side of the petri dish, where the females were later to be released. Thirty-five mung beans, each infested with 2-4 *C. phaseoli* eggs, were spread out in the area outside of the starting zone. Approximately 35 uninfested beans were placed in a small designated circular area (35mm diameter) opposite to the starting zone (**Fig. 1B**). Thus, for females to find the patch with uninfested beans, they first needed to navigate through an area with suboptimal (infested) beans and reject these in favour of potential future hosts of better quality.

Virgin females between 2 and 4 days old were mated to males from the same inbred line. After mating, females were moved individually to a small 35mm petri dish and left there for at least 24 hours without beans (29°C and 12:12LD). Based on a previous study (41), we decided to assay females in groups of five, as this stimulates locomotor activity. A trial started by dividing fifteen mated females equally between three 35mm petri dishes (hereafter: ‘holding dishes’). Arenas with their associated holding dishes were placed on a single heating plate set to 30°C. Females spent five minutes in their holding dish, and approximately one minute before being placed in the arena, an environmental cue was added. To one holding dish, five non-infested seeds were added. We labelled this the “reliable cue”, as it give accurate information about the presence of high-quality, uninfested, future hosts (present at 50% in the arena). To another dish, five highly infested seeds, each with more than 10 eggs, were added. We labelled this the “unreliable cue”, as this treatment provided incorrect information about future host quality (the maximum number of eggs on a seed in the arena was 4, and the average ca. 1.5). To the final dish nothing was added (“no cue” treatment) (**Fig. 1A**). Afterwards, the females were removed from their holding dish and placed together in the starting zone of the arena and left for 10 minutes during which their locations (on uninfested beans or not) were recorded every 30 seconds. At the end of the 10 minutes, the females were placed back into their holding dish for five minutes and the procedure to introduce the environmental cue was repeated. During this time, the uninfested beans in the arena were replaced. The females were exposed to the arena five times (from hereon: “runs”). We predicted that the reliable cue would make females perform more discriminating host choices as these females had gotten correct information about the presence of uninfested, high-quality, seeds. For the no cue and unreliable cue, we expected females to take longer to find the uninfested seeds. For females receiving no cue, we predicted that these females would be less stimulated to start their search for hosts, as they yet had received any information about the presence of hosts, and that they might regard the hosts with competitor eggs acceptable in the absence of other obvious alternatives. For females receiving the unreliable cue, we predicted that these females would spend more time inspecting the infested seeds found in the arena and be more willing to lay eggs on them compared to the other females, as the seeds in the arena are of comparatively good quality (laden with 0-4 eggs) relative to the seeds that the females had been exposed to in the holding dish before each run (laden with > 10 eggs).

The first 12 trials were conducted with two environmental treatments assayed simultaneously (6 trials with reliable versus unreliable cue, and 6 trials with reliable versus no cue assayed in parallel). Once the observer was used to the protocol, we included all three environmental treatments simultaneously, to run an additional 24 trials. The observations were conducted without blinding the observer due to practical limitations. However, to assess the potential for observer-bias, we later repeated the behavioral assays with a blind observer, conducting another 12 trials using the Lome base population. This showed that all three experiments produced qualitatively similar results (**Supplementary 1**). We therefore combined all data for the final analysis presented in the main text, based on 66,000 observations of behavior from 660 females in total.

Adaptive behavioural plasticity (through a learning response) was defined as the improvement between runs (from 1-5) in the rate that females found the patch with uninfested seeds. We first analyzed the number of females found on uninfested beans in each arena as a binomial response using mixed effects generalized linear models using the lme4 package (60) for R (61). We included fixed effects of time (10 minutes within each run) and run (1-5) and their interaction with treatment (reliable/unreliable/no cue). Time and run were both added as continuous variables fitted with a second-degree polynomial. We included the line ID as additional fixed effects. We included the trial ID as a random effect crossed with treatment. We inspected the consistency of the responses in the three lines with reference to treatment and inbreeding status. This showed that the three lines behaved qualitatively the same, and we report a summary of line comparisons in **Supplementary 1**.

We also quantified differences between the three environmental cue treatments by comparing the number of eggs laid on infested and uninfested seeds from the two non-blinded experiments for a total of 480 females scored in fives over 96 trials. One model was run with a binomial response variable to inspect the fraction of eggs laid on the two types of seeds at the end of each trial. This model included environmental cue and line ID as fixed effects. The second model analyzed the number of eggs laid on the uninfested (high-quality) hosts during each of the five consecutive runs as a Poisson distributed response variable. This model included environmental cue, run, and their interaction as fixed effects, and trial as a random term.

### Maternal care and the strength of selection on recessive deleterious alleles

To estimate the fitness consequences of variation in maternal care, we tracked survival of *C. maculatus* larvae competing with *C. phaseoli*. To estimate the strength of selection on recessive deleterious alleles at different levels of competition, we compared survival of inbred and outbred *C. maculatus* larvae. We achieved this by mating parents either within (inbred) or between (outbred) lines (**Fig. 1C**), reasoning that survival differences between inbred and outbred larvae should reflect natural selection against recessive deleterious alleles revealed by homozygosity in inbred individuals (54). The experiment was carried out in two temporal blocks. Two females (mated either within or between lines), were placed in a 35mm petri dish with mung beans. The seeds were either uninfested or infested with a known number (1, 2, 3 or 4) of *C. phaseoli* eggs that had been laid on the seeds a week earlier (as in the behavioural trials). After 72 hours, the females were removed from the petri dishes and the beans were taken out and checked for eggs. If new eggs were found, they were counted, and beans were placed individually into 24-welled chambers. These chambers were placed into climate cabinets set at standard conditions and were later checked to see how many adults that had emerged of each species.

The data consisted of three replicates (each of two females) per line, inbreeding status, and competition level (0-4 *C. phaseoli* eggs), for a total of 60 female duos, laying 8323 eggs on 2450 mung beans. We analysed the survival of *C. maculatus* with a generalized linear mixed effect model using as a binomially distributed response, with inbreeding status (inbred/outbred), the competition intensity, and their interaction as fixed effects. The ID of the female duo was added as a random effect. Note that we chose to sum the eggs of both *C. maculatus* and *C. phaseoli* as a measure of competition intensity, as mung beans had 1-3 *C. maculatus* egg laid on them, with one egg being most common and three eggs rare. Seven seeds had more than three *C. maculatus* eggs on them. We chose to discard these from the analysis since they were unevenly distributed with six of seven seeds from outbred females. A summary of the distribution of eggs laid by both species is presented in **Supplementary 7**.

To predict the strength of selection, we refitted the mixed effect model within a Bayesian framework using the MCMCglmm package (62) for R. Based on the posterior estimates of model coefficients transformed from log-odds back to original scale, we calculated the strength of selection (with 95% credible intervals) for different levels of competition, 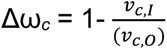, where *v_I_* and *v_O_* is viability of inbred and outbred offspring, respectively. We ran models for 100,000 iterations, preceded by 50,000 burn-in iterations, and stored every 100^th^ iteration, resulting in 1000 uncorrelated posterior estimates of model coefficients. We used flat and weak priors for the random effects.

### Learning and maternal care by mothers of different genetic quality

Using the same set-up as for the original behavioural assays, we quantified the effect of inbreeding load on females’ ability to use the environmental cues to adjust their maternal care behaviour. For this we compared females from within (inbred) and between (outbred) line crosses. To minimize the risk that inbred males’ sperm would negatively affect female fecundity and their motivation for host search, all females were mated with outbred males from the Lome stock population. In each trial, the three cue treatments were included. Nine trials were run with inbred females and nine with outbred females (i.e. three replicates of five females per line and cross type). This resulted in a total of 27,000 behavioural observations from 270 females. The data was analysed by the same model fitting procedure as described above with the addition of adding inbreeding status (inbred/outbred) as a fully crossed fixed effect.

### Genetic load in populations receiving reliable and unreliable environmental information

To quantify the fitness cost associated with the revealed recessive deleterious alleles in each of the three cue treatments, we refitted the mixed models on juvenile survival (see **Fig. 3**) and female behavioural plasticity (see **Fig. 4**) in inbred and outbred individuals using a Bayesian framework. This allowed us to calculate the predicted strength of selection against recessive deleterious alleles based on model posterior distributions. Fitness (*ω*) of inbred and outbred genotypes was calculated by weighting the number of eggs laid on infested and uninfested seeds by the predictions for larval survival at no competition (uninfested seeds) and three competitors (the mean number of *C. phaseoli* eggs on infested seeds in the behavioural experiments). The genome-wide strength of selection against deleterious alleles in treatment *j* was then calculated as: *s_j_* = 1 – *ω_i,j_* / *ω_o,j_.* Using the estimates of *s* we then predicted the long-term fitness consequences and genetic load at mutation-selection balance in environments corresponding to the reliable cue and no cue treatment of our experiment using a previously published model by Kawecki et al. (1997) (13). Details are described in **Supplementary 6**.

## Results

### Learning and maternal care with and without reliable environmental cues

We first explored whether the reliability of environmental cues affected females’ ability to learn the spatial location of uninfested seeds in the arena. Within each of the five runs of a trial, females explored the arena, resulting in more females finding the patch with uninfested seeds as time progressed (Time: 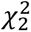 = 4851.9, p < 0.001, **Fig. 2A**). There were clear signs of learning during the trials, signified by a general increase in the proportion of females finding the uninfested hosts between the five consecutive runs (Run: 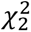 = 5818.0, p < 0.001, **Fig. 2A**). There was evidence for increased adaptive behavioural plasticity in females receiving reliable cues compared to those receiving an unreliable cue or no cue (**Fig. 2A**). This was evident as females receiving the reliable cue finding the uninfested hosts faster (Treatment: 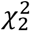 = 108.8, p < 0.001) and showing a faster increase in the behavior between runs (Treatment:Run interaction: 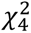 = 72.9, p < 0.001) (**Fig. 2A**). There was also a stronger saturation in the proportion of females found on the uninfested hosts with time within each run in the reliable cue treatment (Treatment:Run:Time interaction: 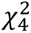 = 116.7, p < 0.001, **Fig. 2A**, **Supplementary Table 2**). Females receiving the reliable cue also laid a larger fraction of eggs (Treatment: 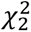 = 204.3, p < 0.001, **Fig. 2B, Supplementary Table 3A**), and more eggs in total (Treatment: 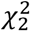 = 224.4, p < 0.001, **Fig. 2B, Supplementary Table 3B**), on uninfested seeds. Thus, reliable environmental cues, providing the opportunity for adaptive behavioural plasticity in maternal care, are likely to reduce competition in larval offspring.

**Figure 2:**
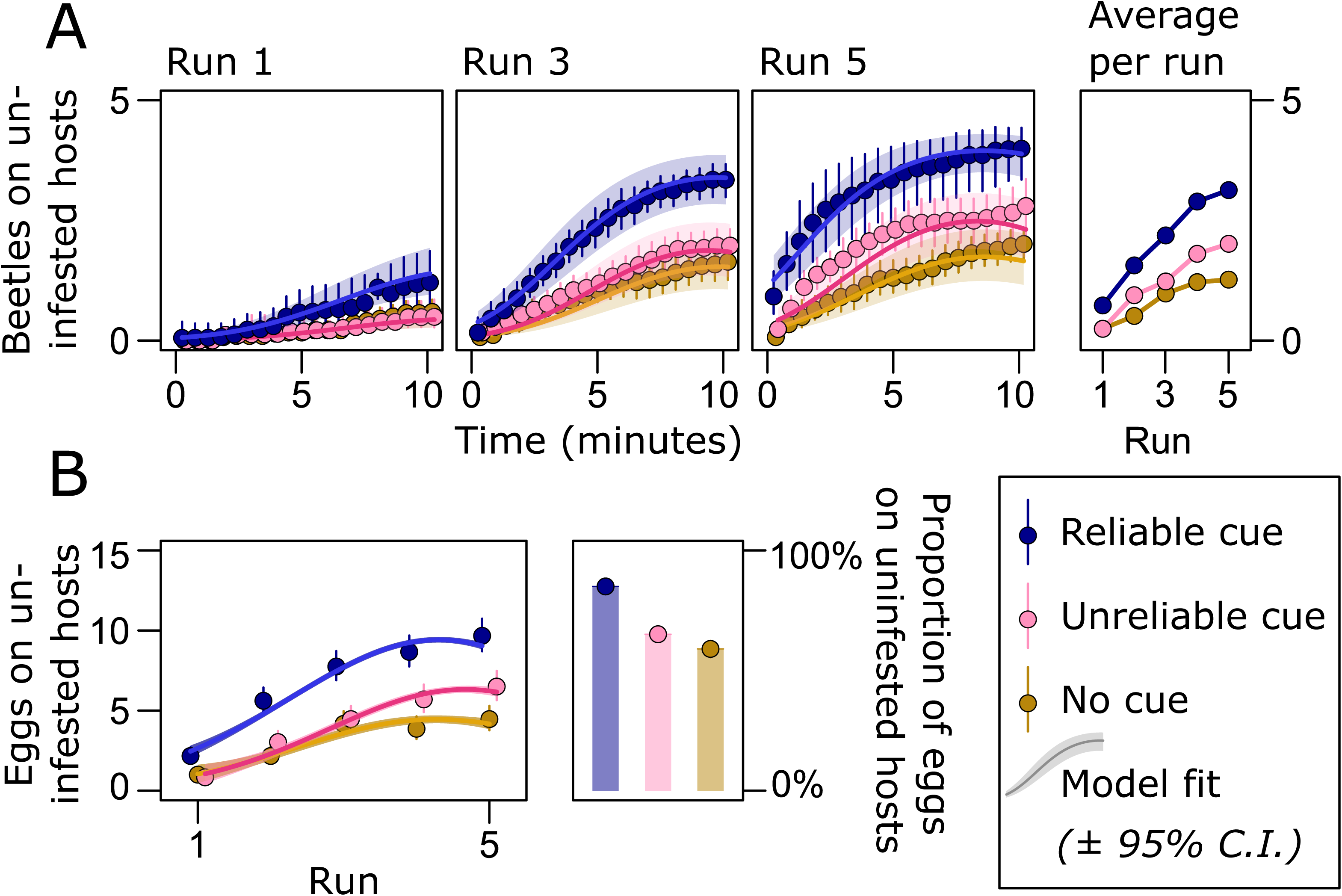
The effect of environmental information on female host search and choice. A) With time, more females located the patch with uninfested host seeds both within and between runs. B) Females also laid more eggs on uninfested hosts in later runs of the experiment. The improvement over runs demonstrates learning during the experiment. Adaptive behavioural plasticity (in form of learning the location of high-quality hosts) was most pronounced in females receiving the reliable cue (blue) compared to those exposed to the unreliable cue (pink) or no cue (brown). This resulted in females receiving the reliable cue placing a higher proportion of their eggs on uninfested seeds relative to females in the other treatments over the entire experiment.

### Maternal care and the strength of selection on recessive deleterious alleles

To estimate the fitness consequences of variation in maternal care, we tracked survival of *C. maculatus* larvae competing with its close relative, *C. phaseoli*. As expected, the density of competitor eggs had a negative impact on egg-to-adult survival of *C. maculatus* (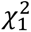 = 52.6, p < 0.001, **Fig 3**). To estimate the strength of selection on recessive deleterious alleles expressed in offspring, we compared the effect of larval competition in inbred and outbred *C. maculatus larvae*. As expected, inbreeding reduced larval survival (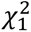 = 49.2, p < 0.001, **Fig. 3**). Moreover, inbred larvae suffered more from the effect of competition compared to outbred offspring (Inbreeding:Infestation intensity interaction: 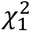 = 3.72, p = 0.035, **Fig. 3**, **Supplementary Table 4**), confirming that competition increases the strength of selection on deleterious alleles.

**Figure 3:**
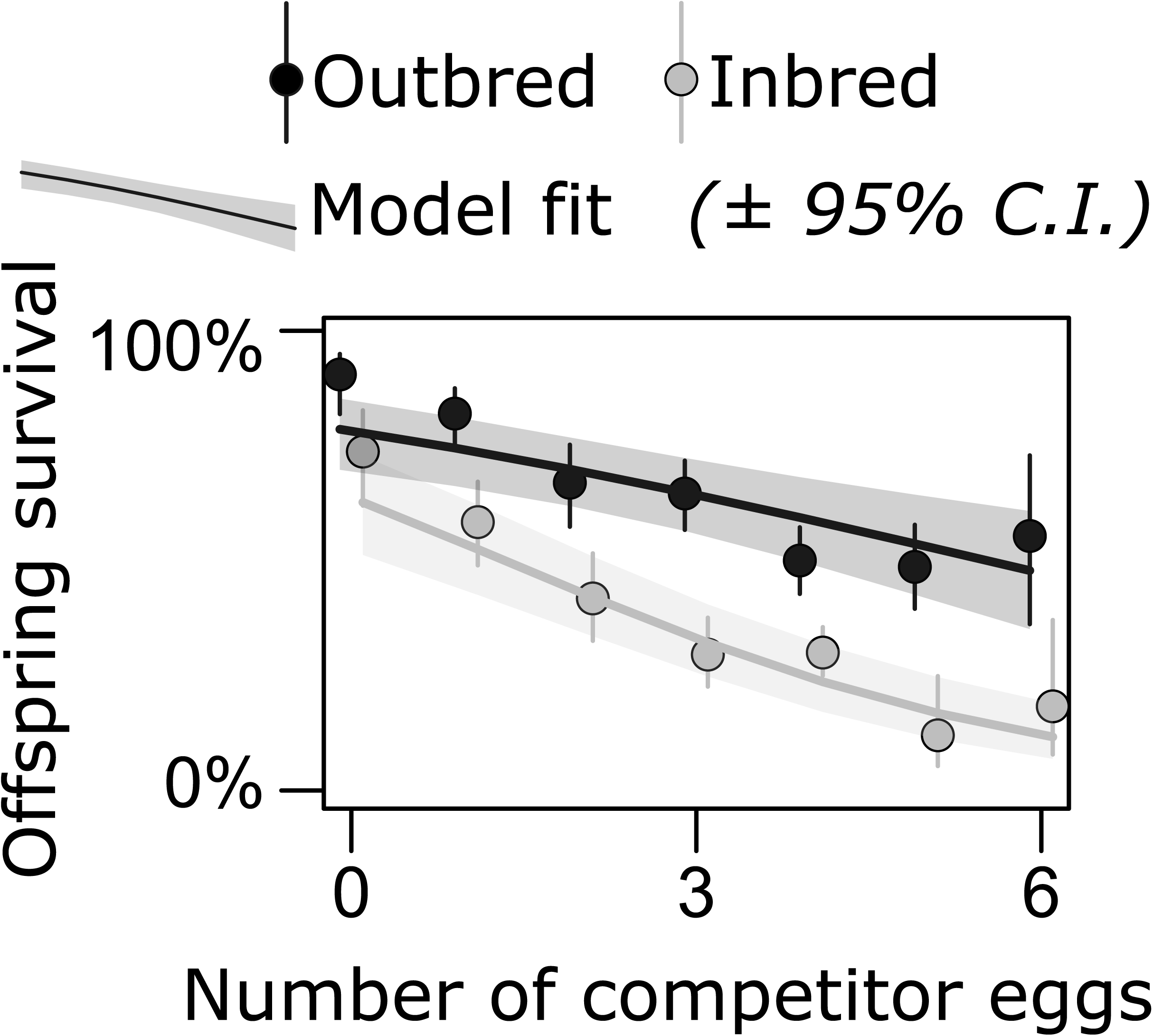
The fitness consequences of larval competition. Eggs placed on seeds with more competitor eggs suffered lower survival. This effect was stronger in inbred (grey) compared to outbred (black) larvae, resulting in stronger selection against deleterious alleles at high competition.

Interaction coefficients in logistic regression models represent deviations from multiplicative effects on odds ratios. However, the relevant currency in the eyes of natural selection is given by relative differences in survival among genotypes. Therefore, we estimated the strength of natural selection against recessive deleterious alleles by comparing the relative survival of outbred and inbred larvae at low density (no competition) and high density (six competitors – the maximum observed within our data range) based on estimates from a Bayesian mixed effect model, with the strength of selection at the level of competition, *c*, proportional to: 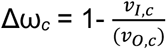, where *v_I_* and *v_O_* is viability of inbred and outbred larvae, respectively. This yielded a fitness cost of deleterious recessives equal to: Δω_C=0_ = 0.20 (95% CI: 0.05-0.37) with no competition, and: Δω_C=6_ = 0.84 (0.67-0.94) at high competition, implying a roughly four times greater strength of purifying selection.

### Learning and maternal care by mothers of different genetic quality

We compared the ability of inbred and outcrossed females to assess the environmental cue and improve maternal care by learning the location of high-quality hosts. A larger proportion of the females receiving the reliable cue were found in the patch with uninfested seeds compared to females in the two other treatments (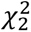 = 177.2, p < 0.001), replicating the results seen in the previous experiment (compare **Fig. 2A** and **Fig. 4A**). Outcrossed females were better at finding the patch with uninfested hosts compared to inbred females (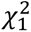 = 16.6, p < 0.001). Interestingly, however, the difference diminished with time both within (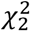 = 117.8, p < 0.001) and between (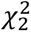 = 223.0, p < 0.001) runs, suggesting that inbred females took longer to find the patch with uninfested seeds, but then performed like outcrossed females in the last run of each trial (**Fig. 4A**). This indicates that learning can diminish innate genetic differences in maternal care. Importantly, this ameliorating effect was strongest in females receiving the reliable cue (Treatment:Inbreeding interaction: 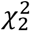 = 14.6, p < 0.001; Treatment:Inbreeding:Run interaction: 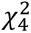 = 24.3, p < 0.001, **Fig. 4A, Supplementary Table 5**). When inspecting the number of eggs laid on host seeds, we found that females receiving the reliable cue laid both a larger fraction of eggs (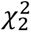 = 72.8, p < 0.001) and overall more eggs (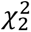 = 98.3, p < 0.001) on uninfested seeds compared to females in the two other treatments, again replicating the results seen in the previous experiment (compare **Fig. 2B** and **Fig. 4B**). Outcrossed females performed better compared to inbred females, both with regards to the fraction (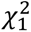 = 13.4, p < 0.001) and total number (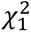 = 33.2, p < 0.001) of eggs laid on uninfested seeds. For the fraction of eggs on uninfested seeds, we found no effect of the environmental cue on the difference between inbred and outcrossed females (Treatment:Inbreeding interaction: 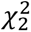 = 3.88, p = 0.14). However, for the total number of eggs on uninfested seeds, the difference between inbred and outcrossed females was smaller in the reliable cue treatment (Treatment:Inbreeding interaction: 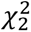 = 11.7, p = 0.003, **Fig. 4B**). This result lends further supports to the hypothesis that reliable environmental cues that leave opportunity for adaptive behavioural plasticity can ameliorate individual fitness differences in maternal care owing to genetic quality.

**Figure 4:**
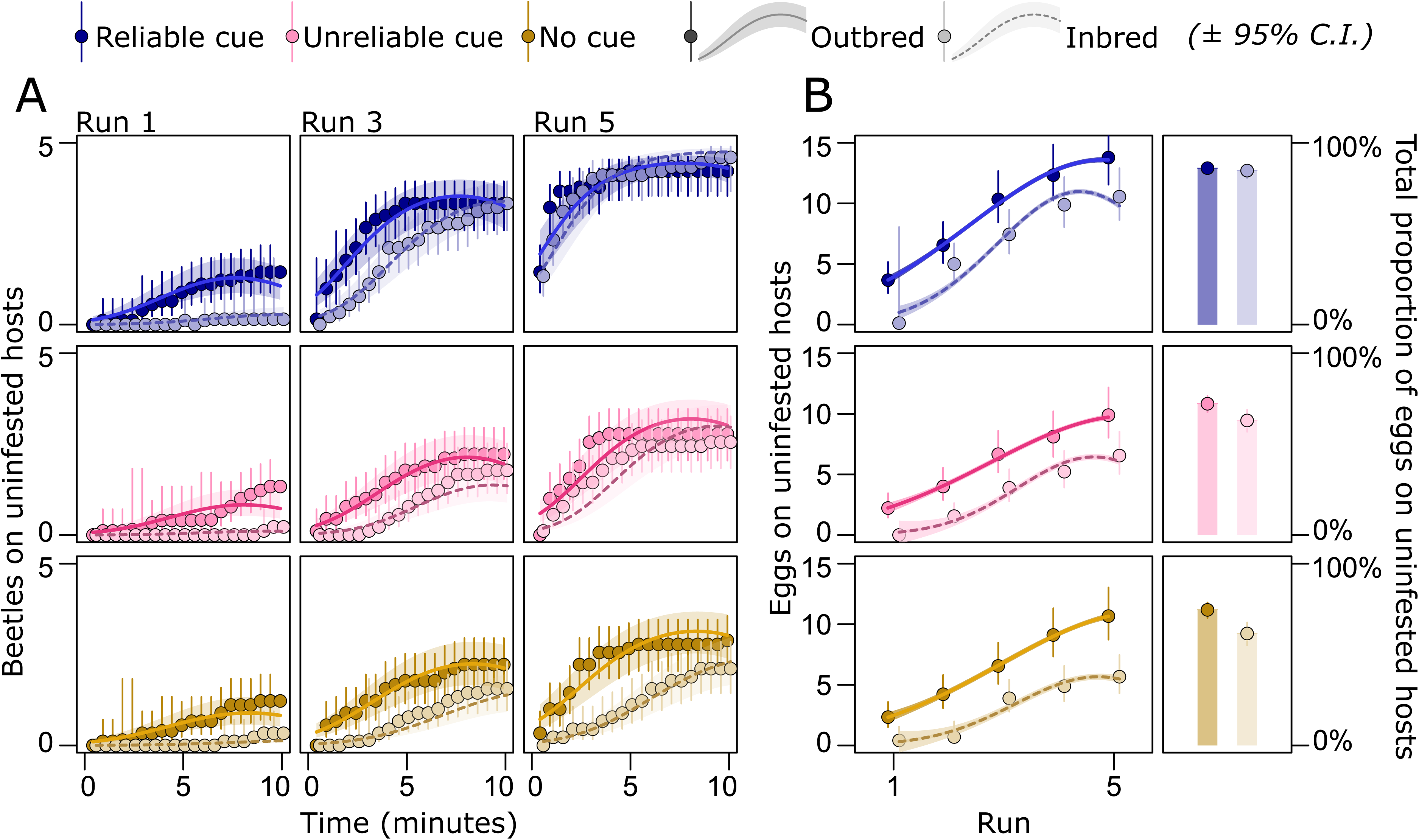
Host search and choice in females of different genetic quality. A) Females learnt the location of the patch with uninfested hosts, and B) laid more eggs on them as runs progressed. The rate of learning was again highest in females exposed to the reliable cue (blue) compared to those exposed to the unreliable cue (pink) or no cue (brown). Inbred females were poorer at finding uninfested seeds. However, adaptive behavioural plasticity (in the form of learning the location of high-quality hosts) reduced this difference between outbred and inbred females, especially if females received the reliable cue. This also resulted in a smaller difference between inbred and outbred females in the proportion of eggs laid on uninfested hosts over the entire experiment.

### Indirect genetic effects depend on environmental information

Combining the information on host choice and juvenile survival across the three genetic lines in inbred and outcrossed state, we estimated the genetic covariance between mean values for maternal care and offspring survival for each genotype and environmental cue treatment. The reliable cue diminished genetic (co)variation between mother and offspring genotype, thus reducing the strength of IGEs relative to the treatments with unreliable cue and no cue (**Fig. 5**).

**Figure 5:**
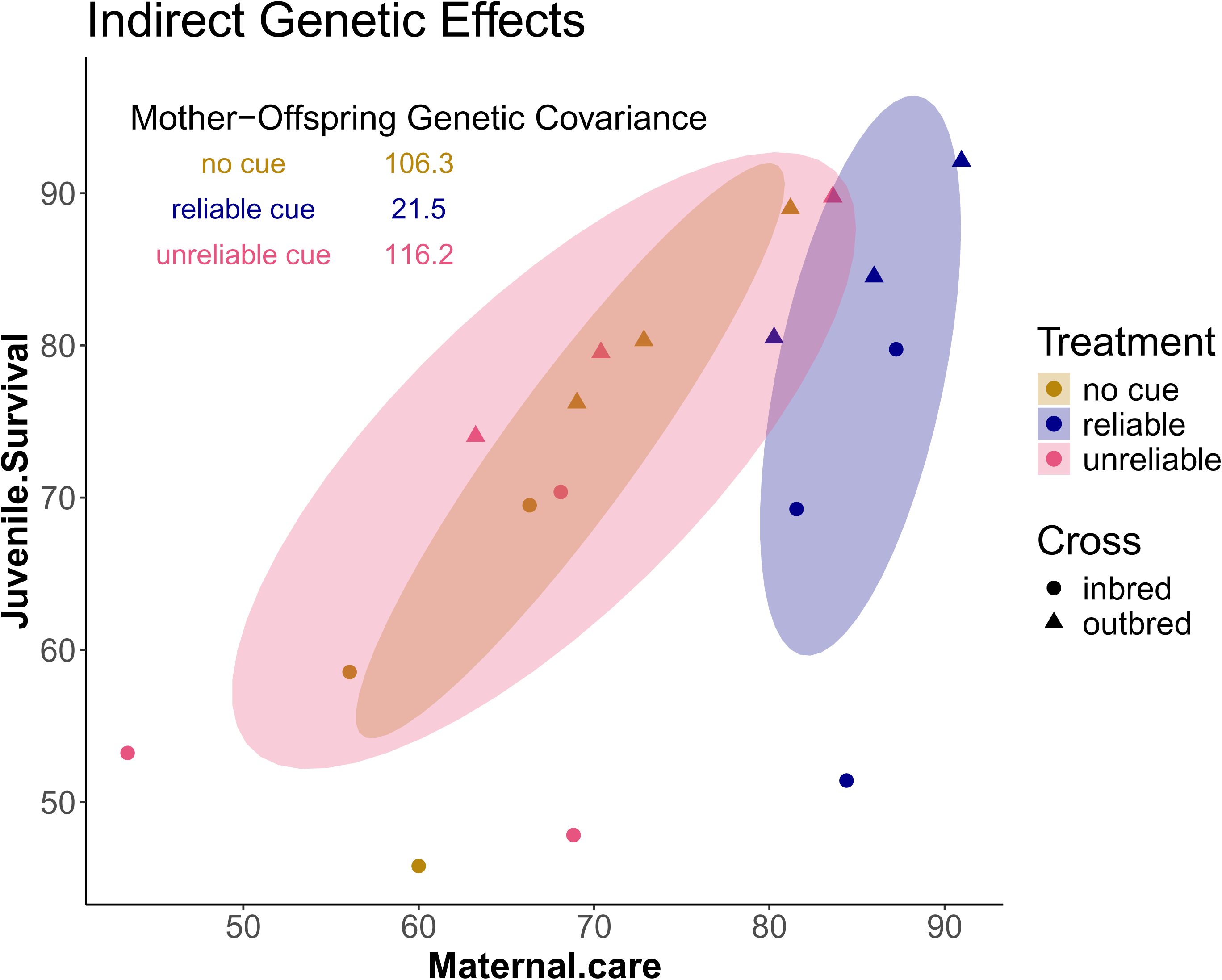
Indirect genetic effects depend on environmental information. Shown are means for each genotype for maternal care (percent eggs laid on uninfested hosts) and juvenile survival (percent expected survival to adulthood) in the treatment with a reliable cue (blue), unreliable cue (pink), and no cue (brown). Outcrossed genotypes are shown as triangles and inbred isogenic lines as circles. Shown are 68% confidence ellipses (corresponding to one standard deviation) based on genotype means.

### Genetic load in populations receiving reliable and unreliable environmental information

We quantified the relative fitness cost associated with recessive deleterious alleles in each of the three treatments by comparing inbred and outbred genotypes. Fitness was calculated by weighting the total number of eggs laid on infested and uninfested seeds (**Fig. 4**) by the predictions for larval survival at no competition (uninfested seeds) and three competitors (the mean number of *C. phaseoli* eggs on infested seeds during behavioural trials) (**Fig. 3**). Expected fitness was higher in the treatment providing reliable cues compared to the other treatments (**Fig. 6A**). Moreover, and echoing the previous findings, the reduction in fitness due to recessive deleterious alleles was lower in the treatment with a reliable cue (Δω= 48%, CI: 29-61%) than in the treatment with an unreliable cue (Δω= 60%, CI: 45-70%, P_MCMC_ = 0.062) and no cue (Δω= 64%, CI: 52-74%, P_MCMC_ = 0.006), although the difference was marginally non-significant in the first comparison (**Fig. 6B**).

**Figure 6:**
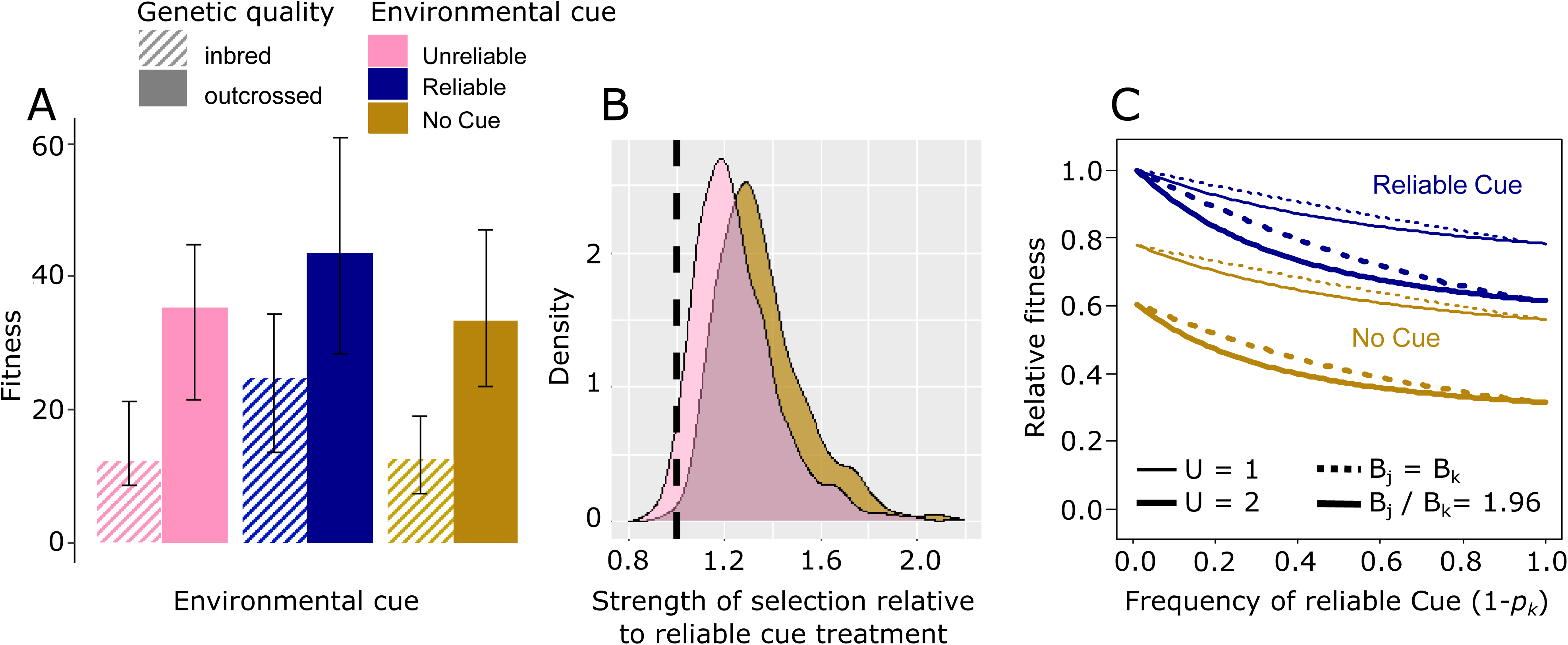
Genetic load in predictable and unpredictable environments. A) Expected offspring production is highest in the treatment providing a reliable cue compared to the other two treatments. B) Bayesian posterior densities showing the increase in the strength of selection in the treatments with unreliable cue and no cue compared to the treatment with a reliable cue. C) Evolution in a predictable environment (corresponding to the reliable cue treatment) is predicted to relax selection on deleterious alleles relative to evolution in an unpredictable environment (corresponding to the no cue treatment), leading to higher fitness but also an accumulation of genetic load in the meta-population. This effect is stronger for higher deleterious mutation rates (thick lines: *U* = 2, thin lines *U* = 1, set to capture the range of probable values in eukaryotes (63)). Load increases further when the predictable environment is more productive (full lines: *B_j_/B_k_* = 1.96 based on the difference in expected fitness of inbred individuals in the treatment with a reliable cue relative to those in the treatment with no cue, and dotted lines: *B_j_* = *B_k_* for comparison). Given that the deleterious mutation rate is not negligible, evolution with adaptive behavioural plasticity in maternal care in the predictable environment will increase load in the unpredictable environment substantially. Details are given in **Supplementary 6**.

Hence, in accordance with theory, reliable environmental information, facilitating adaptive behavioural plasticity, rendered higher fitness and population productivity, while at the same time relaxing selection against deleterious alleles. We predicted the long-term fitness consequences of these dynamics in environments corresponding to our reliable cue (i.e. a predictable environment with accurate information about future conditions available) and no cue treatment (a relatively more unpredictable environment providing no information about future conditions). To do so, we parameterized an existing model by Kawecki et al. (1997) (13) with our treatment-specific estimates of selection and productivity (full details are given in **Supplementary 6**). This model estimates the genetic load at mutation-selection balance in a meta-population experiencing two different environments that differ in the strength of selection on deleterious alleles. Based on our experimental data, we set the mean strength of selection in the unpredictable environment providing no information to be 0.64/0.48 times stronger than in the predictable environment providing reliable information (i.e. *s_k_* = 1.33*s_j_*, **Fig. 6B**). The results how that, while fitness is always greater in the predictable environment with a reliable cue, the frequency of deleterious alleles increases in the population as the environment becomes more predictable due to the alleviating effect of adaptive behavioural plasicity on the strength of selection (**Fig. 6C**). The resulting fitness effects of this accumulated genetic load are more pronounced at high mutation rate and in the unpredictable environment. Thus, in certain scenarios (high reliability of environmental information and high mutation rate), it could be envisioned that species evolving in predictable environments will be unable to colonize unpredictable environments due to competitive exclusion by resident species.

## Discussion

Finetuning of behaviours can allow organisms to avoid perils and shield them from selection. Behavioural plasticity in the form of learning is therefore regarded as a central mechanism allowing animals to cope with fluctuating environments (2,7,21,34). However, by relaxing the strength of selection overall, behavioural plasticity may lead to the accumulation conditionally deleterious alleles, which’ effects may become exposed in the absence of mitigating behaviours. If this process continues for long enough, organisms practising extensive behavioural plasticity are predicted to build up substantial conditionally deleterious genetic loads (9,35,36), perhaps to the extent that they no longer will be competitive in situations where adaptive plasticity is restricted by too rapid and unpredictable environmental changes (63,64). Indeed, to accurately finetune behaviours by learning, the organism needs to be able to assess and predict environmental variability (9,15), leading to the expectation that organisms occupying environments that differ in their predictability may show systematic differences in the expression of learning and conditional genetic loads.

Here, we sought an experimental proof-of-principle for this hypothesis. By manipulating environmental cues given to egg-laying female seed beetles, we show that the benefit of learning is strongly dependent on reliable information of future host quality and location. Reliable environmental cues greatly improved maternal care, in turn reducing genetic selection in juveniles. This relaxed selection is expected to lead to the accumulation of a cryptic genetic load over time. We show that increased genetic load is likely to feed back on the female’s ability to perform maternal care; inbred mothers were significantly worse at finding uninfested hosts. However, inbred females learned the location of uninfested hosts with time, diminishing genetic differences in maternal care. Importantly, the improved learning facilitated by reliable cues contributed further to relaxing selection. Hence, our study presents evidence for a feedback loop between environmental predictability, adaptive behavioural plasticity in form of learning, and the strength of purifying selection.

Our study also provides an experimental demonstration of how the strength of indirect genetic effects (IGEs) can be moulded by the environment. We identified effects of recessive deleterious alleles on both the ability of mothers to provide care and on their offsprings’ survival, such that (outbred) mothers of high genetic quality provided better care and increased survival of their offspring, who also had higher intrinsic viability. This positive genetic covariance between the mothers trait (care) and the expression of her offsprings’ trait (survival) increases genetic variance in offspring fitness relative to a situation without IGEs (34,36,37), and should lead to more efficient purging of genetic load. However, this positive covariance was considerably weakened in the treatment providing reliable cues that reduced genetic differences between both adults and offspring, demonstrating that environmental variation can shape IGEs.

*C. maculatus* females use the presence of eggs to choose whether or not to oviposit (39). However, even though a mother can use cues to make informed decision on where to oviposit, she cannot protect the offspring from the choices made subsequently by other mothers ovipositing on the same host. Likewise, larval genotypes that compete will constitute an evolving environment that may favour the evolution of even more (or less) pronounced competitive traits. This illustrates that environmental predictability in this system is likely to evolve as a function of genes and the IGEs exerted between both competing females and their juvenile offspring (51). Similar dynamics have been elegantly demonstrated in other beetles (12,30,65–67), and as we argue here, the increased opportunity for learning of maternal care may have strong impact on these processes. It is possible that the interplay between environmental variation and IGEs may be a key driver of geographic differentiation in strategies of host use in this system. Indeed, different geographic strains of *C. maculatus* show genetic differentiation in foraging strategies (68) but see: ((69)) and novel host use (45). Host specialization has been shown to evolve rapidly in *C. maculatus* (48,70) and future studies could more directly explore the role of IGEs in driving local adaptation in this system, which may provide important insights into the rapid geographic spread of this insect pest.

While our chosen treatments represent plausible situations for egg-laying *C. maculatus* females, they do not perfectly match environmental variation and frequencies of informed and uninformed oviposition choices in natural populations. Nevertheless, because female *C. maculatus* are faced with ephemeral host availability and varying population densities and levels of competition (39,44,45,45,49,50,71), variation in host quality and abundance is likely ubiquitous. Our treatments were chosen to capture this natural variability, but our results should mainly be taken as a proof-of-principle that environmental variation is key in determining the adaptive value of learning, with cascading effects on the strength of selection and genetic load. To illustrate this point further, we analysed effects on juvenile survival of the competitor *C. phaseoli* when coinhabiting seeds with inbred and outbred *C. maculatus* larvae. This showed that also competing species can be affected by the dynamics between environmental variation and behavioural plasticity (**Supplementary 7**). Indeed, our adoption and parameterization of the model on genetic load in heterogenous environments by Kawecki et al. (1997) (13) illustrates that learning in predictable environments can build up a sizeable conditional genetic load that becomes exposed to selection upon contact with competing species evolving without learning. By comparing an environment giving perfect environmental information with an environment giving no information at all, in a situation where heavily infested hosts were presented before high quality hosts (thus skewing the placement of eggs towards infested hosts), we likely overestimated the cost associated with a learning strategy in the unpredictable environment. Still, it is worth noting that by assuming homogeneous selection across genomic loci within each environment, and by focusing only on deleterious alleles (i.e. no antagonistic pleiotropy across environments), our calculations underestimate the magnitude of the (hidden) genetic load built up in the predictable environment (72).

## Conclusions

To conclude, we find that beetles efficiently provide maternal care when they can learn from previous experience using reliable environmental information. This, in turn, relaxes selection on offspring, allowing for the accumulation of genetic load. We show that the accumulation of load is likely to impact maternal care itself, but that genetic differences in maternal care can be ameliorated by improved learning in environments providing reliable information. The identified eco-evolutionary feedback between learning and the strength of selection is predicted to substantially weaken IGEs between parental provisioning and offspring survival and lead to greater cryptic genetic loads in populations inhabiting predictable environments.

## References

1. Chevin LM, Lande R, Mace GM. Adaptation, Plasticity, and Extinction in a Changing Environment: Towards a Predictive Theory. Kingsolver JG, editor. PLoS Biol [Internet]]. 2010 Apr 27 [cited 2018 Feb 16];8(4):e1000357. Available from: http://dx.plos.org/10.1371/journal.pbio.1000357

2. Price TD, Qvarnström A, Irwin DE. The role of phenotypic plasticity in driving genetic evolution. Proc R Soc Lond B Biol Sci [Internet]]. 2003 Jul 22 [cited 2019 Jan 17];270(1523):1433–40. Available from: http://www.royalsocietypublishing.org/doi/10.1098/rspb.2003.2372

3. Badyaev AV. Stress-induced variation in evolution: from behavioural plasticity to genetic assimilation. Proc R Soc B Biol Sci [Internet]]. 2005 May 7 [cited 2020 Nov 21];272(1566):877–86. Available from: https://royalsocietypublishing.org/doi/full/10.1098/rspb.2004.3045

4. Pfennig DW, editor. Phenotypic Plasticity & Evolution: Causes, Consequences, Controversies [Internet]]. Taylor & Francis; 2021 [cited 2025 Jan 16]. Available from: https://library.oapen.org/handle/20.500.12657/47881

5. Lande R. Adaptation to an extraordinary environment by evolution of phenotypic plasticity and genetic assimilation. J Evol Biol [Internet]]. 2009 [cited 2020 Jan 30];22(7):1435–46. Available from: https://onlinelibrary.wiley.com/doi/abs/10.1111/j.1420-9101.2009.01754.x

6. Snell-Rood EC, Kobiela ME, Sikkink KL, Shephard AM. Mechanisms of Plastic Rescue in Novel Environments. Annu Rev Ecol Evol Syst [Internet]]. 2018 [cited 2019 Dec 27];49(1):331–54. Available from: 10.1146/annurev-ecolsys-110617-062622

7. Huey RB, Hertz PE, Sinervo B. Behavioral Drive versus Behavioral Inertia in Evolution: A Null Model Approach. Am Nat [Internet]]. 2003;161(3):357–66. Available from: http://www.jstor.org/stable/3473156

8. Muñoz MM. The Bogert effect, a factor in evolution. Evolution [Internet]]. 2022 Feb 1 [cited 2024 Feb 8];76(s1):49–66. Available from: 10.1111/evo.14388

9. Snell-Rood EC. An overview of the evolutionary causes and consequences of behavioural plasticity. Anim Behav [Internet]]. 2013 May 1 [cited 2018 Nov 26];85(5):1004–11. Available from: http://www.sciencedirect.com/science/article/pii/S0003347213000080

10. Snell-Rood EC, Van Dyken JD, Cruickshank T, Wade MJ, Moczek AP. Toward a population genetic framework of developmental evolution: the costs, limits, and consequences of phenotypic plasticity. BioEssays [Internet]]. 2010 Jan [cited 2019 Jan 17];32(1):71–81. Available from: http://doi.wiley.com/10.1002/bies.200900132

11. Snell-Rood EC, Burger M, Hutton Q, Moczek AP. Effects of parental care on the accumulation and release of cryptic genetic variation: review of mechanisms and a case study of dung beetles. Evol Ecol [Internet]]. 2016 Apr 1 [cited 2019 Oct 15];30(2):251–65. Available from: 10.1007/s10682-015-9813-4

12. Pascoal S, Shimadzu H, Mashoodh R, Kilner RM. Parental care results in a greater mutation load, for which it is also a phenotypic antidote. Proc R Soc B Biol Sci [Internet]]. 2023 May 24 [cited 2025 Feb 20];290(1999):20230115. Available from: https://royalsocietypublishing.org/doi/10.1098/rspb.2023.0115

13. Kawecki TJ, Barton NH, Fry JD. Mutational collapse of fitness in marginal habitats and the evolution of ecological specialisation. J Evol Biol [Internet]]. 1997 May 1 [cited 2019 Jun 20];10(3):407–29. Available from: https://onlinelibrary.wiley.com/doi/abs/10.1046/j.1420-9101.1997.10030407.x

14. Badyaev AV, Uller T. Parental effects in ecology and evolution: mechanisms, processes and implications. Philos Trans R Soc B Biol Sci [Internet]]. 2009 Apr 27 [cited 2020 Jan 30];364(1520):1169–77. Available from: https://royalsocietypublishing.org/doi/full/10.1098/rstb.2008.0302

15. Gabriel W, Luttbeg B, Sih A, Tollrian R. Environmental Tolerance, Heterogeneity, and the Evolution of Reversible Plastic Responses. Am Nat [Internet]]. 2005 Sep [cited 2025 Feb 21];166(3):339–53. Available from: https://www.journals.uchicago.edu/doi/abs/10.1086/432558

16. Bonamour S, Chevin LM, Charmantier A, Teplitsky C. Phenotypic plasticity in response to climate change: the importance of cue variation. Philos Trans R Soc B Biol Sci [Internet]]. 2019 Mar 18 [cited 2021 Mar 22];374(1768):20180178. Available from: https://royalsocietypublishing.org/doi/10.1098/rstb.2018.0178

17. Vinton AC, Gascoigne SJL, Sepil I, Salguero-Gómez R. Plasticity’s role in adaptive evolution depends on environmental change components. Trends Ecol Evol [Internet]]. 2022 Dec 1 [cited 2024 Sep 30];37(12):1067–78. Available from: https://www.cell.com/trends/ecology-evolution/abstract/S0169-5347(22)00216-6

18. Dupont L, Thierry M, Zinger L, Legrand D, Jacob S. Beyond reaction norms: the temporal dynamics of phenotypic plasticity. Trends Ecol Evol [Internet]]. 2024 Jan 1 [cited 2024 Mar 31];39(1):41–51. Available from: https://www.cell.com/trends/ecology-evolution/abstract/S0169-5347(23)00225-2

19. Dukas R. Evolutionary Biology of Animal Cognition. Annu Rev Ecol Evol Syst [Internet]]. 2004 Dec 15 [cited 2018 Nov 17];35(1):347–74. Available from: http://www.annualreviews.org/doi/10.1146/annurev.ecolsys.35.112202.130152

20. Shettleworth SJ. Animal cognition and animal behaviour. Anim Behav [Internet]]. 2001 Feb 1 [cited 2018 Nov 27];61(2):277–86. Available from]: http://www.sciencedirect.com/science/article/pii/S0003347200916063

21. Baldwin JM. A New Factor in Evolution. Am Nat [Internet]]. 1896 Jun 1 [cited 2018 Dec 21];30(354):441–51. Available from: https://www.journals.uchicago.edu/doi/10.1086/276408

22. Snell-Rood EC, Steck MK. Behaviour shapes environmental variation and selection on learning and plasticity: review of mechanisms and implications. Anim Behav [Internet]]. 2019 Jan 1 [cited 2019 Apr 9];147:147–56. Available from: http://www.sciencedirect.com/science/article/pii/S0003347218302586

23. Caro SM, Griffin AS, Hinde CA, West SA. Unpredictable environments lead to the evolution of parental neglect in birds. Nat Commun [Internet]]. 2016 Mar 29 [cited 2025 Jan 23];7(1):10985. Available from: https://www.nature.com/articles/ncomms10985

24. Fisher DN, Kilgour RJ, Siracusa ER, Foote JR, Hobson EA, Montiglio PO, et al. Anticipated effects of abiotic environmental change on intraspecific social interactions. Biol Rev [Internet]]. 2021 [cited 2025 Feb 27];96(6):2661–93. Available from: https://onlinelibrary.wiley.com/doi/abs/10.1111/brv.12772

25. Sih A. Understanding variation in behavioural responses to human-induced rapid environmental change: a conceptual overview. Anim Behav [Internet]]. 2013 May 1 [cited 2025 Feb 28];85(5):1077–88. Available from: https://www.sciencedirect.com/science/article/pii/S0003347213000936

26. Mousseau T. The adaptive significance of maternal effects. Trends Ecol Evol [Internet]]. 1998 Oct 1 [cited 2019 Jan 17];13(10):403–7. Available from: http://linkinghub.elsevier.com/retrieve/pii/S0169534798014724

27. Clutton-Brock TH. The Evolution of Parental Care. Princeton University Press; 1991. 372 p.

28. Pilakouta N, Jamieson S, Moorad JA, Smiseth PT. Parental care buffers against inbreeding depression in burying beetles. Proc Natl Acad Sci [Internet]]. 2015 Jun 30 [cited 2025 Feb 20];112(26):8031–5. Available from: https://www.pnas.org/doi/full/10.1073/pnas.1500658112

29. Bladon EK, Pascoal S, Kilner RM. Can recent evolutionary history promote resilience to environmental change? Behav Ecol [Internet]]. 2024 Nov 1 [cited 2024 Dec 19];35(6):arae074. Available from: 10.1093/beheco/arae074

30. Mashoodh R, Trowsdale AT, Manica A, Kilner RM. Parental care shapes the evolution of molecular genetic variation. Evol Lett [Internet]]. 2023 Sep 5 [cited 2023 Sep 14];qrad039. Available from: 10.1093/evlett/qrad039

31. Christie ST, Schrater P. Cognitive cost as dynamic allocation of energetic resources. Front Neurosci [Internet]]. 2015 Aug 24 [cited 2025 Feb 28];9:289. Available from: https://www.ncbi.nlm.nih.gov/pmc/articles/PMC4547044/

32. Aiello LC, Wheeler P. The Expensive-Tissue Hypothesis: The Brain and the Digestive System in Human and Primate Evolution. Curr Anthropol [Internet]]. 1995 [cited 2018 Nov 26];36(2):199–221. Available from: https://www.jstor.org/stable/2744104

33. Mery F, Burns JG. Behavioural plasticity: an interaction between evolution and experience. Evol Ecol [Internet]]. 2010 May 1 [cited 2025 Feb 21];24(3):571–83. Available from: 10.1007/s10682-009-9336-y

34. Bailey NW, Marie-Orleach L, Moore AJ. Indirect genetic effects in behavioral ecology: does behavior play a special role in evolution? Simmons L, editor. Behav Ecol [Internet]]. 2018 Jan 13 [cited 2019 Jan 17];29(1):1–11. Available from: https://academic.oup.com/beheco/article/29/1/1/4737155

35. Linksvayer TA, Wade MJ. GENES WITH SOCIAL EFFECTS ARE EXPECTED TO HARBOR MORE SEQUENCE VARIATION WITHIN AND BETWEEN SPECIES. Evolution [Internet]]. 2009 Jul [cited 2019 Jan 17];63(7):1685–96. Available from: http://doi.wiley.com/10.1111/j.1558-5646.2009.00670.x

36. Wolf JB, Iii EDB, Cheverud JM, Moore AJ, Wade MJ. Evolutionary consequences of indirect genetic effects. Trends Ecol Evol [Internet]]. 1998 Feb 1 [cited 2024 Sep 30];13(2):64–9. Available from: https://www.cell.com/trends/ecology-evolution/abstract/S0169-5347(97)01233-0

37. Kirkpatrick M, Lande R. THE EVOLUTION OF MATERNAL CHARACTERS. Evolution [Internet]]. 1989 May 1 [cited 2024 Sep 30];43(3):485–503. Available from: 10.1111/j.1558-5646.1989.tb04247.x

38. Räsänen K, Kruuk LEB. Maternal effects and evolution at ecological time-scales. Funct Ecol [Internet]]. 2007 [cited 2024 Sep 30];21(3):408–21. Available from: https://onlinelibrary.wiley.com/doi/abs/10.1111/j.1365-2435.2007.01246.x

39. Messina FJ, Renwick J a. A. Mechanism of egg recognition by the cowpea weevil Callosobruchus maculatus. Entomol Exp Appl [Internet]]. 1985 [cited 2025 Feb 20];37(3):241–5. Available from: https://onlinelibrary.wiley.com/doi/abs/10.1111/j.1570-7458.1985.tb03481.x

40. Messina FJ, Morrey JL, Mendenhall M. Why do host-deprived seed beetles ?dump? their eggs? Physiol Entomol [Internet]]. 2007 Sep [cited 2019 Jan 17];32(3):259–67. Available from: http://doi.wiley.com/10.1111/j.1365-3032.2007.00579.x

41. Baur J, Nsanzimana J d’Amour, Berger D. Sexual selection and the evolution of male and female cognition: A test using experimental evolution in seed beetles*. Evolution [Internet]]. 2019 [cited 2020 Jan 2];73(12):2390–400. Available from: https://onlinelibrary.wiley.com/doi/abs/10.1111/evo.13793

42. Fox CW. Multiple Mating, Lifetime Fecundity and Female Mortality of the Bruchid Beetle, Callosobruchus maculatus (Coleoptera: Bruchidae). Funct Ecol [Internet]]. 1993 [cited 2018 Feb 19];7(2):203–8. Available from: http://www.jstor.org/stable/2389888

43. Berger D, Stångberg J, Grieshop K, Martinossi-Allibert I, Arnqvist G. Temperature effects on life-history trade-offs, germline maintenance and mutation rate under simulated climate warming. Proc R Soc B]. 2017 Nov 15;284(1866):20171721.

44. Fox CW. A Quantitative Genetic Analysis of Oviposition Preference and Larval Performance on Two Hosts in the Bruchid Beetle, Callosobruchus Maculatus. Evolution [Internet]]. 1993 Feb 1 [cited 2018 Nov 28];47(1):166–75. Available from: https://onlinelibrary.wiley.com/doi/abs/10.1111/j.1558-5646.1993.tb01207.x

45. Messina FJ, Lish AM, Gompert Z. Variable Responses to Novel Hosts by Populations of the Seed Beetle Callosobruchus maculatus (Coleoptera: Chrysomelidae: Bruchinae). Environ Entomol [Internet]]. 2018 Oct 3 [cited 2022 Mar 15];47(5):1194–202. Available from: 10.1093/ee/nvy108

46. Berger D, Stångberg J, Baur J, Walters RJ. Elevated temperature increases genome-wide selection on de novo mutations. Proc R Soc B Biol Sci [Internet]]. 2021 Feb 10 [cited 2022 Jan 5];288(1944):20203094. Available from: https://royalsocietypublishing.org/doi/full/10.1098/rspb.2020.3094

47. Tuda M, Kagoshima K, Toquenaga Y, Arnqvist G. Global Genetic Differentiation in a Cosmopolitan Pest of Stored Beans: Effects of Geography, Host-Plant Usage and Anthropogenic Factors. PLOS ONE [Internet]]. 2014 Sep [cited 2018 Nov 28];9(9):e106268. Available from: https://journals.plos.org/plosone/article?id=10.1371/journal.pone.0106268

48. Fricke C, Arnqvist G. RAPID ADAPTATION TO A NOVEL HOST IN A SEED BEETLE (*CALLOSOBRUCHUS MACULATUS*): THE ROLE OF SEXUAL SELECTION: ADAPTATION AND SEXUAL SELECTION. Evolution [Internet]]. 2007 Feb [cited 2018 Feb 19];61(2):440–54. Available from: http://doi.wiley.com/10.1111/j.1558-5646.2007.00038.x

49. Mitchell R. The Evolution of Oviposition Tactics in the Bean Weevil, Callosobruchus maculatus (F.). Ecology [Internet]]. 1975 May 1 [cited 2018 Nov 28];56(3):696–702. Available from: https://esajournals.onlinelibrary.wiley.com/doi/abs/10.2307/1935504

50. Credland PF, Dick KM, Wright AW. Relationships between larval density, adult size and egg production in the cowpea seed beetle, Callosobruchus maculatus. Ecol Entomol [Internet]]. 1986 Feb 1 [cited 2018 Nov 28];11(1):41–50. Available from: https://onlinelibrary.wiley.com/doi/abs/10.1111/j.1365-2311.1986.tb00278.x

51. Fox CW, Savalli UM. Inheritance of Environmental Variation in Body Size: Superparasitism of Seeds Affects Progeny and Grandprogeny Body Size Via a Nongenetic Maternal Effect. Evolution [Internet]]. 1998 [cited 2024 Sep 30];52(1):172–82. Available from: https://onlinelibrary.wiley.com/doi/abs/10.1111/j.1558-5646.1998.tb05150.x

52. Berger D, Grieshop K, Lind MI, Goenaga J, Maklakov AA, Arnqvist G. INTRALOCUS SEXUAL CONFLICT AND ENVIRONMENTAL STRESS: SEX, GENES, AND CONFLICT IN STRESSFUL ENVIRONMENTS. Evolution [Internet]]. 2014 May [cited 2018 Feb 19];n/a-n/a. Available from: http://doi.wiley.com/10.1111/evo.12439

53. Grieshop K, Berger D, Arnqvist G. Male-benefit sexually antagonistic genotypes show elevated vulnerability to inbreeding. BMC Evol Biol [Internet]]. 2017 Dec [cited 2018 Nov 28];17(1). Available from: http://bmcevolbiol.biomedcentral.com/articles/10.1186/s12862-017-0981-4

54. Charlesworth D, Willis JH. The genetics of inbreeding depression. Nat Rev Genet [Internet]]. 2009 Nov [cited 2020 Dec 5];10(11):783–96. Available from: https://www.nature.com/articles/nrg2664

55. Grieshop K, Maurizio PL, Arnqvist G, Berger D. Selection in males purges the mutation load on female fitness. Evol Lett [Internet]]. 2021 [cited 2022 Mar 10];5(4):328–43. Available from: https://onlinelibrary.wiley.com/doi/abs/10.1002/evl3.239

56. Temreshev II, Kazenas VL. Callosobruchus phaseoli (Gyllenhal, 1833) (Coleoptera, Chrysomelidae, Bruchinae): a new invasive species in Kazakhstan. Acta Biol Sib [Internet]]. 2020 Jul 23 [cited 2024 Oct 15];6:87–92. Available from: https://abs.pensoft.net/article/53070/

57. Lambrides CJ, Imrie BC. Susceptibility of mungbean varieties to the bruchid species Callosobruchus maculatus (F.), C. phaseoli (Gyll.), C. chinensis (L.), and Acanthoscelides obtectus (Say.) (Coleoptera:Chrysomelidae). Aust J Agric Res [Internet]]. 2000 [cited 2024 Oct 15];51(1):85–90. Available from: https://www.publish.csiro.au/ar/ar99051

58. Arnqvist G, Rönn J, Watson C, Goenaga J, Immonen E. Concerted evolution of metabolic rate, economics of mating, ecology, and pace of life across seed beetles. Proc Natl Acad Sci [Internet]]. 2022 Aug 16 [cited 2024 Jun 12];119(33):e2205564119. Available from: https://www.pnas.org/doi/abs/10.1073/pnas.2205564119

59. Bagchi B, Corbel Q, Khan I, Payne E, Banerji D, Liljestrand-Rönn J, et al. Sexual conflict drives micro- and macroevolution of sexual dimorphism in immunity. BMC Biol [Internet]]. 2021 Jun 2 [cited 2021 Jun 4];19(1):114. Available from: 10.1186/s12915-021-01049-6

60. Bates D. Mixed models in R using the lme4 package Part 4: Inference based on profiled deviance].:23.

61. R Core Team. R: A Language and Environment for Statistical Computing [Internet]. Vienna, Austria: R Foundation for Statistical Computing]; 2020. Available from: https://www.R-project.org/

62. Hadfield JD. MCMC methods for multi-response generalized linear mixed models: the MCMCglmm R package. J Stat Softw]. 2010;33(2):1–22.

63. Agrawal AF, Whitlock MC. Mutation Load: The Fitness of Individuals in Populations Where Deleterious Alleles Are Abundant. Annu Rev Ecol Evol Syst [Internet]]. 2012 Dec [cited 2018 Feb 19];43(1):115–35. Available from: http://www.annualreviews.org/doi/10.1146/annurev-ecolsys-110411-160257

64. Tienderen PHV. Evolution of Generalists and Specialists in Spatially Heterogeneous Environments. Evolution [Internet]]. 1991 [cited 2021 Mar 22];45(6):1317–31. Available from: https://onlinelibrary.wiley.com/doi/abs/10.1111/j.1558-5646.1991.tb02638.x

65. Rohner PT, Moczek AP. Vertically inherited microbiota and environment modifying behaviours conceal genetic variation in dung beetle life history. Proc R Soc B Biol Sci [Internet]]. 2024 Apr 17 [cited 2025 Jun 26];291(2021):20240122. Available from: https://royalsocietypublishing.org/doi/10.1098/rspb.2024.0122

66. Bladon EK, Pascoal S, Bird N, Mashoodh R, Kilner RM. The evolutionary demise of a social interaction: experimentally induced loss of traits involved in the supply and demand of care. Evol Lett [Internet]]. 2023 Jun 1 [cited 2025 Mar 4];7(3):168–75. Available from: 10.1093/evlett/qrad016

67. Rohner PT, Jones JA, Moczek AP. Plasticity, symbionts and niche construction interact in shaping dung beetle development and evolution. J Exp Biol [Internet]]. 2024 Mar 7 [cited 2025 Jun 26];227(Suppl_1):jeb245976. Available from: 10.1242/jeb.245976

68. Credland PF, Dick KM. Food consumption by larvae of three strains of Callosobruchus maculatus (Coleoptera: Bruchidae). J Stored Prod Res [Internet]]. 1987 Feb [cited 2022 Mar 15];23(1):31–40. Available from: https://linkinghub.elsevier.com/retrieve/pii/0022474X87900336

69. Burc E, Girard-Tercieux C, Metz M, Cazaux E, Baur J, Koppik M, et al. Life-history adaptation under climate warming magnifies the agricultural footprint of a cosmopolitan insect pest. Nat Commun [Internet]]. 2025 Jan 18 [cited 2025 Jan 19];16(1):827. Available from: https://www.nature.com/articles/s41467-025-56177-2

70. Rêgo A, Messina FJ, Gompert Z. Dynamics of genomic change during evolutionary rescue in the seed beetle Callosobruchus maculatus. Mol Ecol]. 2019 May;28(9):2136–54.

71. Utida S. Density dependent polymorphism in the adult of Callosobruchus maculatus (Coleoptera, Bruchidae). J Stored Prod Res [Internet]]. 1972 Jun 1 [cited 2018 Nov 28];8(2):111–25. Available from: http://www.sciencedirect.com/science/article/pii/0022474X72900288

72. Hermisson J, Wagner GP. The Population Genetic Theory of Hidden Variation and Genetic Robustness. Genetics [Internet]]. 2004 Dec [cited 2018 Jan 29];168(4):2271–84. Available from: http://www.genetics.org/lookup/doi/10.1534/genetics.104.029173

