## Supplementary Material 1-7 for "The reliability of environmental cues shape learning and selection against deleterious alleles in seed beetles"

*Article doi:* 10.1098/rspb.xxxx.xxxx

#### INDEX:

**Supplementary 1:** Repeatability across genetic lines and experiments

**Supplementary 2:** Spatial learning and finding uninfested hosts

**Supplementary 3:** Effects of environmental cues and learning on the number of eggs laid on infested and uninfested hosts.

**Supplementary 4:** Offspring fitness consequences

**Supplementary 5:** Maternal care and learning in females different genetic quality

**Supplementary 6**: Genetic load in populations receiving reliable and unreliable environmental information

**Supplementary 7:** Effects on competitive exclusion

### Supplementary 1: Repeatability across genetic lines and experiments

*Table S1a: Females found in the patch with uninfested host seeds for each line*

| Line | Run | No Cue | Reliable Cue | Unreliable Cue |
|---|---|---|---|---|
| L15:11 | 1 | 0.26 | 0.40 | 0.04 |
| L15:12 | 2 | 0.41 | 1.10 | 0.59 |
| L15:13 | 3 | 0.52 | 1.26 | 0.68 |
| L15:14 | 4 | 0.79 | 1.99 | 1.29 |
| L15:15 | 5 | 0.92 | 2.68 | 1.19 |
| L3:10 | 1 | 0.22 | 0.91 | 0.33 |
| L3:10 | 2 | 0.62 | 1.77 | 0.80 |
| L3:10 | 3 | 1.11 | 2.63 | 1.13 |
| L3:10 | 4 | 1.11 | 3.76 | 1.57 |
| L3:10 | 5 | 1.10 | 4.05 | 1.72 |
| L35:15 | 1 | 0.23 | 0.77 | 0.13 |
| L35:15 | 2 | 0.53 | 1.86 | 0.96 |
| L35:15 | 3 | 1.30 | 2.76 | 2.08 |
| L35:15 | 4 | 1.58 | 3.52 | 2.87 |
| L35:15 | 5 | 1.64 | 4.33 | 3.58 |

*Table S1b: Females found in the patch with uninfested host seeds in each experiment*

| Line | Data | Run | No Cue | Reliable Cue | Unreliable Cue |
|---|---|---|---|---|---|
| all | 2-plate | 1 | 0.50 | 1.48 | 0.53 |
| all | 2-plate | 2 | 0.78 | 2.20 | 1.39 |
| all | 2-plate | 3 | 1.20 | 2.34 | 2.02 |
| all | 2-plate | 4 | 1.08 | 2.85 | 2.71 |
| all | 2-plate | 5 | 0.73 | 3.26 | 3.03 |
| all | 3-plate | 1 | 0.17 | 0.30 | 0.08 |
| all | 3-plate | 2 | 0.45 | 1.26 | 0.63 |
| all | 3-plate | 3 | 0.91 | 2.15 | 1.11 |
| all | 3-plate | 4 | 1.18 | 3.20 | 1.71 |
| all | 3-plate | 5 | 1.34 | 3.90 | 1.94 |
| Lome base | blinded | 1 | 0.28 | 0.85 | 0.44 |
| Lome base | blinded | 2 | 0.50 | 1.53 | 1.36 |
| Lome base | blinded | 3 | 1.00 | 2.10 | 1.07 |
| Lome base | blinded | 4 | 1.36 | 2.31 | 1.54 |
| Lome base | blinded | 5 | 1.40 | 1.46 | 1.65 |

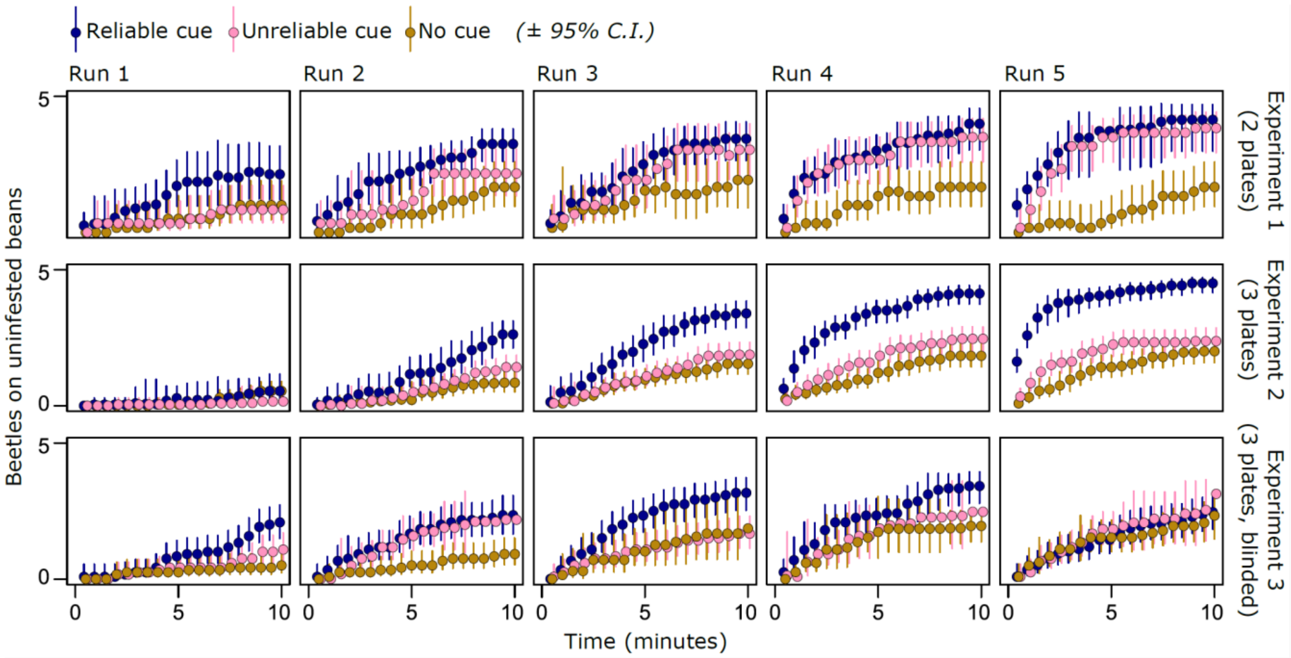


**Supplementary Figure S1a: Female host search in the three separate experiments**
Experiment 1 was run using only 2 treatments/plates in a trial, whereas experiments 2 and 3 were run with all three treatments simultaneously. Experiment 3 was done with a blind observed to confirm the previous results and check for observer bias. However, experiment 3 used older females from the Lome base population that had already laid eggs, which may explain the saturating effect in runs 4 and 5 for females receiving the reliable cue. There was a significant effect of cue treatment in all three experiments when analyzed separately. Experiment 2 contains almost three times more data than the other two experiments, but all data was analyzed together for the analyses in the main manuscript.

| *Table S1c: Egg laying of each line* | | | | |
|---|---|---|---|---|
| **Line** | **trait** | **No Cue** | **Reliable Cue** | **Unreliable Cue** |
| L15:11 | prop. Eggs on uninfested | 0.46 | 0.79 | 0.52 |
| L3:10 | prop. Eggs on uninfested | 0.60 | 0.87 | 0.62 |
| L35:15 | prop. Eggs on uninfested | 0.62 | 0.86 | 0.71 |
| L15:11 | Total eggs | 17.8 | 29.6 | 23.1 |
| L3:10 | Total eggs | 25.3 | 39.1 | 22.7 |
| L35:15 | Total eggs | 36.4 | 50.8 | 48.8 |

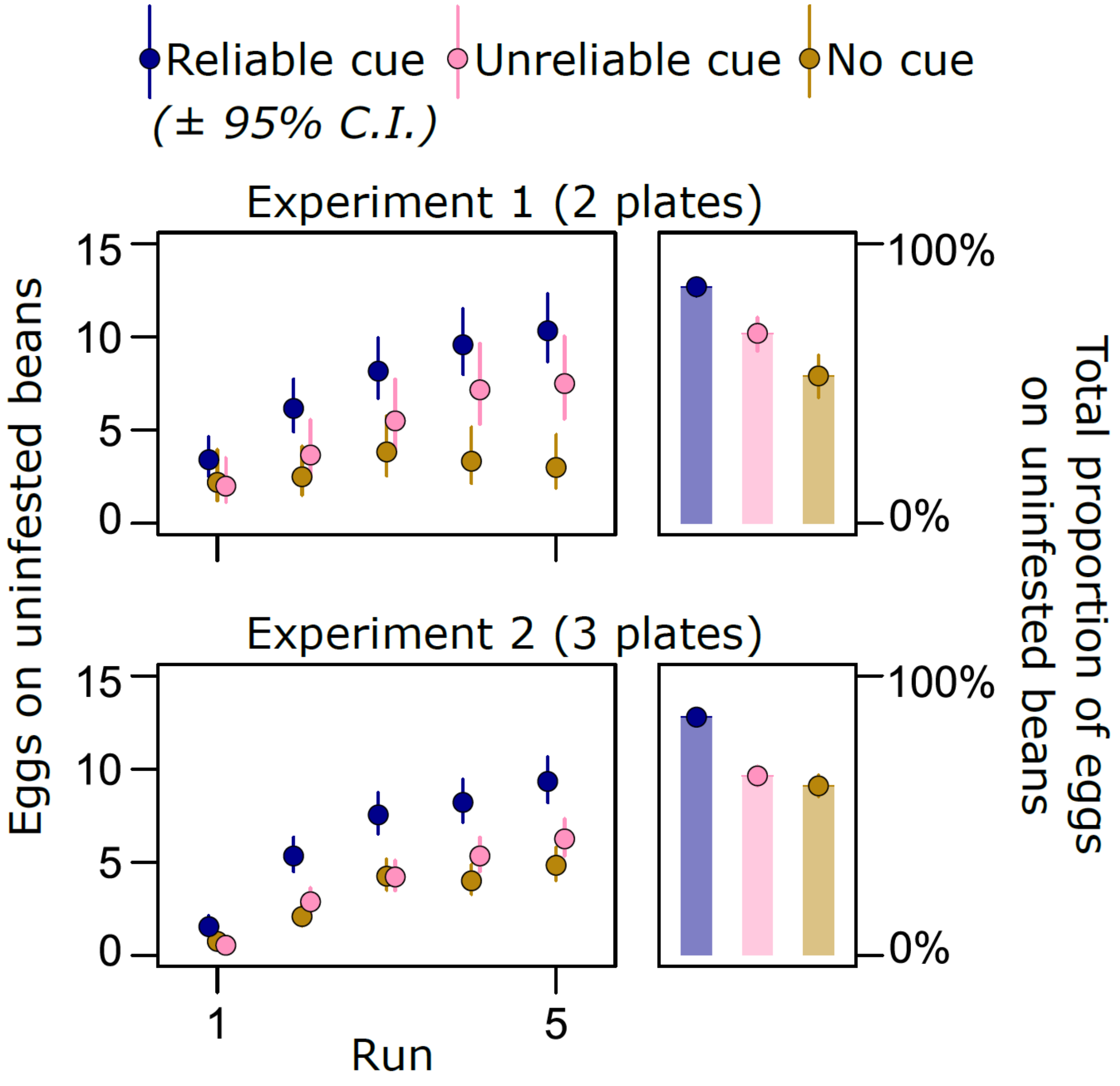


**Supplementary Figure S1b: Female host choice in the two separate experiments**
Experiment 1 was run using only 2 treatments/plates in a trial, whereas experiment 2 was run with all three treatments simultaneously. Experiment 2 contains almost three times more data than the experiment 1, but all data was analyzed together for the analyses in the main manuscript.

*Table S1d: Juvenile survival of each line*

| Line | Competitors | inbred | outbred |
|---|---|---|---|
| L15:11 | 0 | 0.83 | 0.94 |
| L15:11 | 1 | 0.55 | 0.88 |
| L15:11 | 2 | 0.37 | 0.60 |
| L15:11 | 3 | 0.39 | 0.70 |
| L15:11 | 4 | 0.31 | 0.48 |
| L15:11 | 5 | 0.22 | 0.57 |
| L15:11 | 6 | 0.13 | 0.47 |
| L3:10 | 0 | 0.54 | 0.87 |
| L3:10 | 1 | 0.47 | 0.76 |
| L3:10 | 2 | 0.41 | 0.52 |
| L3:10 | 3 | 0.21 | 0.52 |
| L3:10 | 4 | 0.29 | 0.54 |
| L3:10 | 5 | 0.22 | 0.35 |
| L3:10 | 6 | NA | 0.63 |
| L35:15 | 0 | 0.83 | 0.88 |
| L35:15 | 1 | 0.73 | 0.73 |
| L35:15 | 2 | 0.44 | 0.67 |
| L35:15 | 3 | 0.39 | 0.63 |
| L35:15 | 4 | 0.31 | 0.44 |
| L35:15 | 5 | 0.14 | 0.50 |
| L35:15 | 6 | 0.25 | 0.67 |

### Supplementary 2: spatial learning and finding uninfested hosts

**Supplementary Table 2:** The proportion of females found on uninfested beans throughout trials

```
     AIC      BIC   logLik deviance df.resid
 26350.6  26530.3 -13151.3  26302.6    13176

Scaled residuals:
    Min      1Q  Median      3Q     Max
-3.5635 -0.5612 -0.1912  0.5308 15.0752

Random effects:
 Groups     Name        Variance Std.Dev.
 Cue:Trial2 (Intercept) 0.649    0.8056
 Trial2     (Intercept) 0.405    0.6364
Number of obs: 13200, groups:  Cue:Trial2, 132; Trial2, 48
Analysis of Deviance Table (Type II Wald chisquare tests)

Response: cbind(Good, Bad)
                                           Chisq Df Pr(>Chisq)
Cue                                     108.8032  2  < 2.2e-16 ***
scale(poly(Run, 2))                    5817.9877  2  < 2.2e-16 ***
scale(poly(Time, 2))                   4851.8744  2  < 2.2e-16 ***
Line                                     21.4320  3  8.562e-05 ***
Cue:scale(poly(Run, 2))                  72.9073  4  5.519e-15 ***
Cue:scale(poly(Time, 2))                  6.4015  4     0.1711
scale(poly(Run, 2)):scale(poly(Time, 2)) 116.7392  4  < 2.2e-16 ***
```

#### **Supplementary 3:** Effects of environmental cues and learning on the number of eggs laid on infested and uninfested hosts.

**Supplementary Table 3A:** Fraction of eggs laid on uninfested host seeds.

```
     AIC      BIC   logLik deviance df.resid
   445.8    461.2   -216.9    433.8       90

Scaled residuals:
     Min       1Q   Median       3Q      Max
-2.40319 -0.57555  0.00169  0.59550  1.60384

Random effects:
 Groups Name        Variance Std.Dev.
 Trial2 (Intercept) 0.005627 0.07501
Number of obs: 96, groups:  Trial2, 36

Analysis of Deviance Table (Type II Wald chisquare tests)

Response: cbind(tot.eggs.good, eggs.inf)
       Chisq Df Pr(>Chisq)
Cue   204.302  2  < 2.2e-16 ***
Line   36.592  2  1.133e-08 ***
```

**Supplementary Table 3B:** Number of eggs laid on uninfested host seeds

```
     AIC      BIC   logLik deviance df.resid
  2113.8   2163.8  -1044.9   2089.8      467

Scaled residuals:
    Min      1Q  Median      3Q     Max
-3.0645 -0.8069 -0.0370  0.5533  6.6224

Random effects:
 Groups Name        Variance Std.Dev.
 Trial2 (Intercept) 0.05269  0.2295
Number of obs: 479, groups:  Trial2, 36

Analysis of Deviance Table (Type II Wald chisquare tests)

Response: eggs.good
                   Chisq Df Pr(>Chisq)
Cue             224.3792  2  < 2.2e-16 ***
poly(Run, 2)    366.0223  2  < 2.2e-16 ***
Line             50.6002  2  1.029e-11 ***
Cue:poly(Run, 2)  6.9067  4     0.1409
```

### Supplementary 4: Offspring fitness consequences

**Supplementary Table 4:** Larval survival of inbred and outbred offspring as a function of number of competitors

```
     AIC      BIC   logLik deviance df.resid 
  3592.3   3638.8  -1788.2   3576.3     2439 

Scaled residuals: 
    Min      1Q  Median      3Q     Max 
-3.6293 -0.7961  0.2800  0.7451  3.1662 

Random effects:
 Groups    Name        Variance Std.Dev.
 Female.ID (Intercept) 0.5713   0.7559  
Number of obs: 2447, groups:  Female.ID, 90

Analysis of Deviance Table (Type II Wald chisquare tests)

Response: cbind(emerge.Cm, eggs.Cm - emerge.Cm)
                     Chisq Df Pr(>Chisq)    
inbred             49.2025  1  2.309e-12 ***
competitors        52.6453  1  3.996e-13 ***
line                3.7208  2    0.15561    
experiment          0.0036  1    0.95246    
inbred:competitors  4.4337  1    0.03524 *  
```

#### Supplementary 5: maternal care and learning in females different genetic quality

**Supplementary Table 5A**: Fraction of inbred and outbred females on uninfested beans

```
Random effects:
 Groups       Name        Variance Std.Dev.
 Beans:Trial2 (Intercept) 0.1568   0.3960
 Trial2       (Intercept) 0.1715   0.4141
Number of obs: 5400, groups:  Beans:Trial2, 54; Trial2, 18
```

```
 Analysis of Deviance Table (Type II Wald chisquare tests)

 Response: cbind(GB, W)
                              Chisq Df Pr(>Chisq)
 Beans                     177.1784  2  < 2.2e-16 ***
 poly(Run, 2)             2789.1718  2  < 2.2e-16 ***
 poly(Time, 2)            1984.3466  2  < 2.2e-16 ***
 inbred                     16.6210  1  4.564e-05 ***
 Line                        8.5895  2  0.0136403 *
 Beans:poly(Run, 2)        134.9163  4  < 2.2e-16 ***
 Beans:poly(Time, 2)         5.1680  4  0.2704872
 Beans:inbred               14.6004  2  0.0006754 ***
 poly(Run, 2):inbred       222.9991  2  < 2.2e-16 ***
 poly(Time, 2):inbred      117.8378  2  < 2.2e-16 ***
 Beans:poly(Run, 2):inbred  24.3108  4  6.920e-05 ***
 Beans:poly(Time, 2):inbred  2.4440  4  0.6546889
```

**Supplementary Table 5B:** Fraction of eggs laid on uninfested beans by inbred and outbred females

```
Random effects:
 Groups Name        Variance Std.Dev.
 Trial2 (Intercept) 0        0
Number of obs: 54, groups:  Trial2, 18
```

```
Analysis of Deviance Table (Type II Wald chisquare tests)

Response: cbind(eggs.tot.good, eggs.inf)
               Chisq Df Pr(>Chisq)
Beans        72.8049  2  < 2.2e-16 ***
inbred       13.4449  1  0.0002457 ***
Line          5.7064  2  0.0576607 .
Beans:inbred  3.8833  2  0.1434699
inbred:Line  28.0956  2  7.927e-07 ***
```

**Supplementary Table 5C:** Number of eggs laid on uninfested beans by inbred and outbred females throughout runs of the trials

```
     AIC      BIC   logLik deviance df.resid
  1098.7   1174.3   -528.4   1056.7      249

Scaled residuals:
    Min      1Q  Median      3Q     Max
-2.3084 -0.4943 -0.0433  0.3601  3.7731

Random effects:
 Groups Name        Variance Std.Dev.
 Trial2 (Intercept) 0.02096  0.1448
Number of obs: 270, groups:  Trial2, 18
```

```
Analysis of Deviance Table (Type II Wald chisquare te

Response: eggs.good
                             Chisq Df Pr(>Chisq)
Beans                      98.3300  2  < 2.2e-16 ***
inbred                     33.1878  1  8.367e-09 ***
poly(Run, 2)              304.5710  2  < 2.2e-16 ***
Line                       11.2131  2   0.003674 **
Beans:inbred               11.6582  2   0.002941 **
Beans:poly(Run, 2)          4.4384  4   0.349919
inbred:poly(Run, 2)        28.0330  2  8.179e-07 ***
Beans:inbred:poly(Run, 2)   2.0173  4   0.732575
```

#### Supplementary 6: **Genetic load in populations receiving reliable and unreliable environmental information**

Based on our experimental estimates of selection, we predicted the long-term fitness consequences of behavioural plasticity in the predictable (reliable cue) and unpredictable (no cue) environment. Following Haldane's (1937) original derivation [1], for an unconditionally deleterious allele at mutation-selection balance, the genetic load at locus *i* is: $L_i \approx 2qs_i$, where q is the frequency of the deleterious allele, $s$ is the strength of purifying selection, and the factor of 2 accounts for diploidy. Considering only deleterious alleles and assuming strong selection so that genetic drift is negligible and $q$ is rare, $q_{eq} \approx \mu_i/s_i$ at mutation-selection balance, and thus, $L_i \approx 2s_i\mu_i/s_i = 2\mu_i$. This yields Haldane's original, somewhat surprising, result indicating that in a constant environment the fitness cost of segregating deleterious alleles is independent of the strength of selection against them. Assuming multiplicative fitness effects across loci, fitness is then [1]:

$$W \approx \prod_i^1 (1 - s_i \frac{2\mu_i}{s_i}) \approx e^{-U} \qquad \text{[Eq. 1]}$$

where $U$ is the diploid genome-wide deleterious mutation rate. However, this result does not hold when environments vary, because the equilibrium frequency of deleterious alleles is a result of the genome-wide mean strength of selection across variable conditions, whereas the fitness load at any point in time is determined by the strength of selection in the current environment [2,3]. Following [4], we set environment $j$ as our predictable environment with a reliable cue, and $k$ as our unpredictable environment with no cue, and calculate the expected fitness in each environment in a meta-population encountering them at frequency $p_j$ and $p_k$:

$$W_j = \prod_i^1 (1 - \frac{2\mu_i}{c_j + c_k s_{k,i}/s_{j,i}}) \qquad \text{[Eq. 2a]}$$

$$W_k = \prod_i^1 (1 - \frac{2\mu_i}{c_j s_{j,i}/s_{k,i} + c_k}) \qquad \text{[Eq. 2b]}$$

where $c_j$ and $c_k$ is the relative contribution to total population growth from environment $j$ and $k$, respectively, given by:

$$c_k = \frac{\phi_k \prod_i^1 (1 - 2q_i s_{k,i})}{\prod_i^1 (1 - 2q_i s_{j,i}) + \phi_k \prod_i^1 (1 - 2q_i s_{k,i})} = \frac{\phi_k W_k}{W_j + \phi_k W_k} \qquad \text{[Eq. 3a]}$$

$$c_j = 1 - c_k \qquad \text{[Eq. 3b]}$$

with $\phi_k = B_k p_k / B_j p_j$, where $B_k$ and $B_j$ is the productivity in environment $k$ and $j$, respectively. Based on our experimental data, we set $B_k$ and $B_j$ equal to the estimated mean offspring production in each environment (Fig. 6A, main text), and the mean strength of selection in the unpredictable environment to be 0.64/0.48 times stronger in the unpredictable environment (i.e. $s_k = 1.33s_j$, Fig. 6B, main text). We then used equations 2 and 3 to calculate the expected fitness in the predictable and unpredictable environment at mutation-selection balance for different mutation rates and values of $p_k$ and $p_j$ (1-$p_k$). For simplicity, we assumed constant mutation rates across loci, and that selection acts with the same strengths across all loci within a given environment, noting that variability in *s* and μ will tend to further increase the effects reported [5]. Results are reported in the main text.

#### References


1. Haldane, J. B. S. The Effect of Variation of Fitness. *Am. Nat.* **71**, 337–349 (1937).
2. Kawecki, T. J. Adaptation to Marginal Habitats. *Annu. Rev. Ecol. Evol. Syst.* **39**, 321–342 (2008).
3. Agrawal, A. F. & Whitlock, M. C. Mutation Load: The Fitness of Individuals in Populations Where Deleterious Alleles Are Abundant. *Annu. Rev. Ecol. Evol. Syst.* **43**, 115–135 (2012).
4. Kawecki, T. J., Barton, N. H. & Fry, J. D. Mutational collapse of fitness in marginal habitats and the evolution of ecological specialisation. *J. Evol. Biol.* **10**, 407–429 (1997).
5. Hermisson, J. & Wagner, G. P. The Population Genetic Theory of Hidden Variation and Genetic Robustness. *Genetics* **168**, 2271–2284 (2004).

#### Supplementary 7: Effects on competitive exclusion

To further explore potential consequences of the build-up of genetic load for the outcome of interspecific competition, we analysed how the survival of the competitor, *C. phaseoli*, depended on the density of eggs and the inbreeding status of *C. maculatus* in the experiment on larval survival (Fig. 3, main text). We also estimated the competitive outcome between the two species by comparing the relative numbers of adults emerging from coinhabited beans. Both models thus fitted survival as a binomial response as a function of the number of *C. phaseoli* and *C. maculatus* eggs, with the latter variable crossed with inbreeding status of *C. maculatus*. The ID of the female duo that laid the *C. maculatus* eggs was added as a random effect in both models. To assess the predicted outcome of interspecific competition depending on the genetic load of *C. maculatus*, we calculated interspecific competition coefficients between the species, separately for inbred and outbred *C. maculatus*, based on how each species' larval survival was affected by competing eggs of its own and the other species. We approximated each species' carrying capacity for a single mung bean based on the survival data (K ≈ 2 for both species). Based on these approximations, we calculated interspecific competition isoclines and mapped them on the model predictions of relative survival of the two species.

Survival of *C. phaseoli* depended on if *C. maculatus* is inbred or not (Inbreeding:Infestation intensity interaction: $\chi^2_1$ = 5.38, p = 0.020). Likewise, the outcome of the species-interaction depended on the inbreeding status of *C. maculatus* in such a way that *C. phaseoli* has a competitive disadvantage against outbred *C. maculatus* larvae, but an advantage when *C. macultatus* is inbred (Inbreeding:Infestation intensity: $\chi^2_1$ = 7.83, p = 0.005, Fig. S7a). Based on estimates of interspecific competition coefficients and carrying capacities for each species based on the survival data, we calculated competition isoclines and mapped them on the model predictions of competitive outcome between the two species. When *C. phaseoli* is competing against outbred *C. maculatus*, a non-stable equilibrium is reached, with the more abundant species excluding the rare colonizing species. However, when *C. phaseoli* is competing against inbred *C. maculatus* it is the superior competitor and is predicted to exclude *C. maculatus*, irrespective of starting population sizes (Fig. S7a).

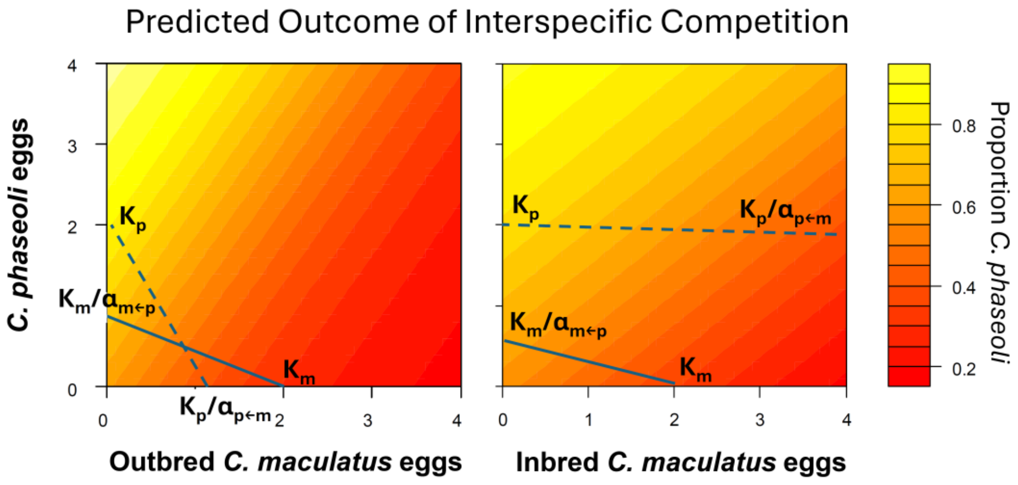


**Supplementary Fig. S7a: Model predictions for larval survival of *C. phaseoli* as a function of the number of *C. maculatus* eggs and the genetic load of *C. maculatus*.**
The relative success of *C. phaseoli* in the survival experiment when competing against either outcrossed or inbred *C. maculatus*. Interspecific competition isoclines are drawn based on approximations of each species' carrying capacity (K=2) and interspecific competition coefficients (α) on a single mung bean, based on the survival data. When *C. phaseoli* is competing against outbred *C. maculatus*, a non-stable equilibrium is predicted, with the competitive outcome dependent on the starting abundance of each species. Thus, the more abundant species is expected to exclude the colonizing species. When *C. phaseoli* is competing against inbred *C. maculatus* it is the superior competitor and will exclude *C. maculatus*, irrespective of starting frequencies.

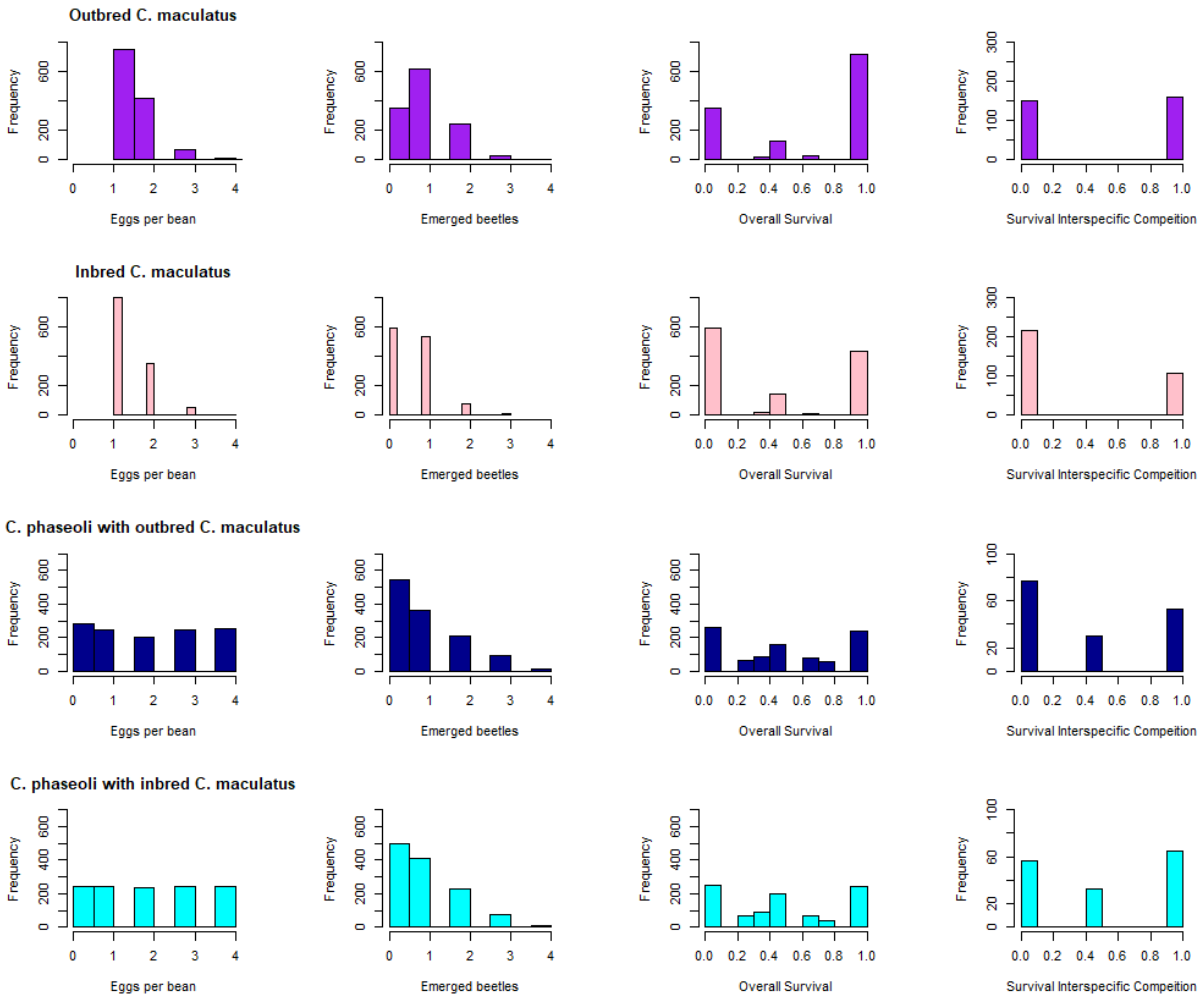


**Supplementary Fig. S7b: Eggs laid and beetles emerged for *C. maculatus* and *C. phaseoli* in the juvenile competition experiment.**

Furthest left column; distributions of eggs laid per bean for inbred and outbred *C. maculatus*, and for *C. phaseoli* competing against inbred and outbred *C. maculatus*. Note that beans from *C. phaseoli* were picked so that a uniform density distribution was attained for the experiment. Second column; the number of emerged adult beetles from single seeds. Third column the distributions of juvenile survival from single seeds. Fourth column; the distributions of juvenile survival isolating effects of competition: for C. maculatus (top two rows), data is based on seeds with only a single *C. maculatus* egg, but more than two *C. phaseoli* eggs; for *C. phaseoli* (bottom two rows) the data is based on seeds with one or two *C. phaseoli* eggs and at least two *C. maculatus* eggs. Note that *C. maculatus* are slightly bigger than C. phaseoli.
